# Adjusting ADJUST: Optimizing the ADJUST Algorithm for Pediatric Data Using Geodesic Nets

**DOI:** 10.1101/753822

**Authors:** Stephanie C. Leach, Santiago Morales, Maureen E. Bowers, George A. Buzzell, Ranjan Debnath, Daniel Beall, Nathan A. Fox

**Affiliations:** University of Maryland, College Park, MD

**Keywords:** Electroencephalography, EEG artifacts, automated artifact classification algorithm, developmental research, geodesic sensor net, independent component analysis

## Abstract

A major challenge for electroencephalograph (EEG) studies on pediatric populations is that large amounts of data are lost due to artifacts (e.g., movement and blinks). Independent component analysis (ICA) can separate artifactual and neural activity, allowing researchers to remove such artifactual activity and retain a greater percentage of EEG data for analyses. However, manual identification of artifactual components is time consuming and requires subjective judgment. Automated algorithms, like ADJUST and ICLabel, have been validated on adults using the international 10-20 system, but to our knowledge no such algorithms have been optimized for the geodesic sensor net, which is often used with infants and children. Therefore, in an attempt to automate artifact selection for pediatric data collected with geodesic nets, we modified ADJUST’s algorithm. Our “adjusted-ADJUST” algorithm was compared to the “original-ADJUST” algorithm and ICLabel in adults, children, and infants on three different performance measures: respective classification agreement with expert coders, the number of trials retained following artifact removal, and the reliability of the EEG signal after pre-processing with each algorithm. Overall, the adjusted-ADJUST algorithm performed better than the original-ADJUST algorithm and no ICA correction with adult and pediatric data. Moreover, it performed better than ICLabel for pediatric data. These results indicate that optimizing existing algorithms for data collected with a geodesic net improves artifact classification and retains more trials, potentially facilitating EEG studies with pediatric populations.

## 1. Introduction

Electroencephalography (EEG) is a useful, non-invasive method for studying brain functioning (Luck, 2014). EEG is acquired by placing electrodes or sensors on the scalp to record electrical signals generated by the brain. However, in addition to recording neural activity, these sensors also record environmental noise, such as muscle activity from movements and changes of the electrical dipoles of the eyes during blinks or saccades (Luck, 2014). These non-neural sources of activity are often referred to as artifacts, and they can contaminate signals of interest in the EEG and potentially lead to the misinterpretation of results (Luck, 2014). This problem is exacerbated in pediatric research because children often have a hard time sitting still and focusing for long periods of time. As a result, studies involving children are often shorter (incorporating fewer trials) and a greater proportion of trials must ultimately be rejected due to artifact contamination, as compared to adult studies. As such, preprocessing methods that can remove artifacts instead of deleting whole trials are of special importance in developmental research.

One such artifact removal method is independent component analysis (ICA), which decomposes the EEG data into maximally independent components, separating artifacts from neural data (for more detail on how ICA works as an artifact rejection method see Urigüen & Garcia-Zapirain, 2015 or Jung, Makeig, Westerfield, Townsend, Courchesne, & Sejnowski, 2000). Once components reflecting artifacts are identified, they can be removed and the data can be reconstructed, thereby subtracting artifacts from the EEG data without having to remove entire epochs of data (Jung et al., 2000). However, the identification of artifactual components takes time and often requires subjective judgment. Moreover, the number of components generated by ICA depends on the number of unique sources of information (i.e., electrodes present in the dataset), such that the number of components is often equal to the number of electrodes used for recording (Luck, 2014). For high-density channel systems, which can have 64, 128, or even 256 channels, manually inspecting each component to determine if it is artifactual can be quite time consuming. Furthermore, while artifacts generally show consistent activation patterns (e.g., high activity over frontal sites and low activity over posterior sites for blinks; Chaumon, Bishop, & Busch, 2015), there are many cases where it is unclear whether a component reflects an artifact or neural activity that follows a similar activation pattern. As such, determining whether or not an ambiguous component reflects neural or artifactual activity is not always straightforward and requires subjective judgement. Relying on subjective judgements, which have the potential for introducing systematic bias into the data, can be especially problematic for large, multi-site studies that rely on multiple people to analyze the data. That is because not all individuals will select the same ICA components for removal and they may unintentionally remove components that reflect neural data, which can affect Event-Related Potentials (ERPs) in a variety of ways (e.g., increasing/decreasing component amplitude). For these reasons, using subjective judgement to identify components reflecting artifactual activity can impair attempts to replicate findings of independent research groups. That is, removing or retaining a different subset of ICA components could potentially be the difference between a significant and a non-significant effect.

In response to these issues, numerous automated artifact selection algorithms have been created to expedite the artifact selection process and increase objectivity. One such widely used algorithm is ADJUST (Mognon, Jovicich, Bruzzone, & Buiatti, 2011), which uses a combination of the spatial and temporal features of the components to classify them as artifactual or neural. In our experience, ADJUST often misclassifies components in pediatric data collected with geodesic nets (e.g., components reflecting blinks are not classified as artifactual), which may result from the fact that ADJUST was validated using adult data collected with a 10-20 system. Pediatric samples often use geodesic nets because they are easier and faster to apply compared to gel-based systems. Due to their shorter attention spans, children are more likely to lose interest in completing EEG tasks if capping takes a long time. Similarly, with infants, the longer capping takes the more likely the infant is to get upset, which will result in the collection of noisy EEG data during the task; in order to allow children and infants to complete more trials, capping should happen as quickly as possible (Brooker,et al., 2019). As such, geodesic nets are quite popular in developmental research. In order to facilitate research in pediatric populations, an automated artifact selection algorithm that is optimized for data collected with geodesic nets is needed.

In addition to ADJUST, a recently released algorithm, called ICLabel (Pion-Tonachini, Kreutz-Delgado, & Makeig, 2019), provides another automated artifact selection method by using a machine learning classifier. The validation paper for ICLabel (Pion-Tonachini et al., 2019) also presents analyses suggesting that ADJUST’s performance is lower than previously thought, even for adult data collected via the 10-20 system. While ICLabel exhibited superior performance on adult data, the authors noted that there was anecdotal evidence suggesting underperformance with infant data that was likely due to a lack of infant data in the training dataset (ICLabel’s machine-learning approach is optimized based on a labeled set of training data). Moreover, even though there were a few studies with children included in the training dataset for ICLabel, the majority of the studies in the training dataset were with adult populations using the 10-20 system. Ultimately, ICLabel, like ADJUST, is optimized for adults and the 10-20 system, which means that it may also underperform on pediatric data collected with geodesic nets. In fact, to the best of our knowledge, the performance of these algorithms on pediatric data has not been formally evaluated and no automated component classification algorithms have been optimized for pediatric data collected with geodesic nets.

To rectify this, we have made modifications to the ADJUST algorithm in an attempt to optimize component classification for pediatric data collected with geodesic nets. More specifically, we modified the ocular artifact selection algorithms to use the spatial layout of a geodesic net and we added an algorithm that ensures selected artifacts do not also include evidence of neural activity, such as a local maximum within the alpha band. In order to examine the performance of our updated algorithm, which we refer to as “adjusted-ADJUST”, we tested the classification accuracy of our adjusted-ADJUST algorithm, as well as the *original* ADJUST algorithm (“original-ADJUST”) and ICLabel algorithms. We further examined EEG data quality following the application of each of these algorithms to pediatric and adult EEG data sets. Specifically, we tested these algorithms by measuring their respective classification agreement with expert coders, the number of trials retained following artifact removal, and the reliability of the EEG signal after pre-processing with each algorithm. We predicted that component classifications using the adjusted-ADJUST algorithm would better match manual classifications by experienced coders as compared to the original-ADJUST algorithm and ICLabel for pediatric data, whereas differences would likely not emerge for adult data. Moreover, we expected that for pediatric but not adult data, the adjusted-ADJUST algorithm (as compared to the original-ADJUST, ICLabel, and no ICA correction at all) would also retain more trials and yield greater internal consistency reliability.

## 2. Methods

### 2.1 Test Data Sets

#### 2.1.1 Data Set 1

The adjusted-ADJUST algorithm was tested on an existing dataset that contained both child and adult data so that the algorithm’s performance on the two populations could be compared. This test data set consisted of 10 children (4 males, *M*=7.9 years, *SD* = 0.8 years, range 7.2 to 9.75 years) and 10 adults (3 males, *M*=21.6 years, *SD*=0.6 years, range 20.5 to 22.6) who participated in a larger study examining brain activity associated with action observation and action execution. The University of Maryland Institutional Review Board approved of all the study protocols. Before participating in the study, all adult participants gave written informed consent while all child participants and their parents gave written informed assent and consent, respectively.

Participants performed an action execution and action observation task. In this task, participants watched three different videos with a jittered onset time. Each trial started with a fixation cross that was present for 500 ms to 1500 ms, followed by one of the three videos containing two boxes. The three videos corresponded to one of three conditions: Action Observation, Action Execution, and Scene Observation (control). In the Action Observation condition, the participant watched a hand grasp one of the boxes. For the Action Execution condition, the participant executed a grasping action while watching a similar video containing boxes. Finally, the Scene Observation condition contained similar background stimuli as the other two conditions, but it did not depict a grasping action, nor did it require the participant to execute a grasping action. Because the stimuli across conditions were almost identical, we expected no differences across conditions in early visual ERP components and there were no differences in mean P1 ERP amplitude between any of the three video conditions, F(2,36)=0.102,p=0.903. As such, we did not consider video condition as a factor in any of the later analyses. There were 40 trials in each condition, with a total of 120 trials overall. More details on the task can be found in Morales, Bowman, Velnoskey, Fox, and Redcay (2019).

#### 2.1.2 Data Set 2

The adjusted-ADJUST algorithm was also tested on an existing infant dataset so that the algorithm’s performance on infant data could be assessed. This test data set consisted of 10 infants (5 males, *M*=5.2 months, *SD* = 0.6 months, range 4.2 to 6 months) who participated in a larger study examining temperament and brain development. The University of Maryland Institutional Review Board approved of all the study protocols. Before participating in the study, all participant parents provided written informed consent.

Participants completed a resting state task. In order to keep the infants attention and prevent excessive motor movement, an experimenter showed the infants different colored balls and turned a bingo wheel. The resting state task was divided into six, 30-second blocks. This resulted in a total of 180, one-second segments for analysis. For more details about the experimental procedure see Fox, Henderson, Rubin, Calkins, and Schmidt (2001).

#### 2.1.3 EEG Preprocessing

Continuous EEG data were collected for both data sets using a 128-channel HydroCel Geodesic Sensor Net and EGI Netstation software (version 4; Electrical Geodesic Inc., Eugene, OR). Consistent with recommended acquisition protocols for high-impedance EEG systems (Electrical Geodesic Inc., Eugene, OR), the target impedance level for the electrodes was below 50kΩ during data collection. The EEG signal was amplified through an EGI NetAmps 300 amplifier with a sampling rate of 500 Hz and referenced online to the vertex electrode (Cz). Data were then preprocessed using the EEGLAB toolbox (Delorme & Makeig, 2004) and custom MATLAB scripts (The MathWorks, Natick, MA). Because the infant data files ranged from four to eight minutes in length, the number of electrodes was reduced from 128 to 64 in order to ensure a good ICA decomposition for all participants. Channels were deleted such that the layout matched that of a 64-channel HydroCel Geodesic Sensor net. The data were high-pass filtered at 0.3 Hz and low-pass filtered at 49 Hz. Bad channels were identified and removed globally using the FASTER toolbox (Nolan, Whelan, & Reilly, 2010); On average, 4.4 channels were removed by FASTER, with the number of removed channels ranging from one to eight. This dataset was then copied and put through a 1.0 Hz high-pass filter before being epoched into arbitrary one-second segments. Any epochs where channel voltage exceeded 1000 μV or power within the 20-40 Hz band (after Fourier analysis) exceeded 30 dB were deleted. Alternatively, if a particular channel caused more than 20% of the epochs to be marked for rejection, that channel was rejected instead of the epochs. After running ICA on this copied dataset, any components that were marked for rejection, using either adjusted-ADJUST (which is the focus of the ensuing methods sections), original-ADJUST, or ICLabel, were marked for rejection. Then, the weights from the copied dataset were subsequently applied back to the original dataset (see Viola, Debener, Thorne, & Schneider, 2010, for further details on this approach) and the components that were marked for rejection were removed. Data were then epoched into 1500 millisecond segments that started 500 milliseconds before the video onset. After ICA artifact removal and epoching, a two-step procedure for identifying residual artifacts was employed. First, any epochs where ocular channel (E1, E8, E14, E21, E25, or E32 for dataset 1; E1, E9, E22, or E32 for dataset 2) voltages exceeded ±125 μV, indicating the presence of residual ocular activity not removed through ICA, were rejected. Second, for any epoch in which non-ocular channel voltages exceeded ±125 μV, these channels were interpolated at the epoch level, unless greater than 10% of the channels exceeded ±125 μV, in which case the epoch was rejected instead. Finally, all missing channels were interpolated using a spherical spline interpolation and then the data were referenced to the average of all the electrodes. The average number of interpolated channels per epoch for the first (128-channel) data set (including those globally rejected) was 4.8 for adjusted-ADJUST, 4.0 for original-ADJUST, 4.6 for ICLabel, and 3.0 when not using any form of ICA correction. For the second (64-channel) data set, the average number of interpolated channels per epoch (including those globally rejected) was 1.3 for adjusted-ADJUST, 1.1 for original-ADJUST, 1.2 for ICLabel, and 1.0 when not using any form of ICA correction. The number of interpolated channels per epoch for the first (128-channel) data set ranged from one to 19 for both adjusted-ADJUST and ICLabel, one to 18 for original-ADJUST, and one to 20 when not using any form of ICA correction. For the second (64-channel) data set, the number of interpolated channels per epoch ranged from one to 12 for adjusted-ADJUST and ICLabel and one to 11 for original-ADJUST and no form of ICA correction.

### 2.2 Changes to the ADJUST Algorithm

Similar to Mognon and colleagues (2011), normalized component topographies were used when computing spatial information. The original-ADJUST classified artifacts into four different categories: eye blinks, horizontal eye movements, vertical eye movements, and discontinuities (e.g., pop-offs). Various functions using temporal or spatial information were used to calculate the likelihood that a component belonged to one of the four artifact classes. In order to improve ADJUST performance on pediatric data, changes were made to the eye blink detection and horizontal eye movement detection functions. The vertical eye movement and generic discontinuity detection functions were not altered because anecdotal evidence suggested that these functions already showed high performance on data collected with geodesic nets. Additionally, new lines of code were added that check for local maxima within the alpha band for all components that were classified as artifacts. For the adjusted-ADJUST code, see the online supplement [Upon publication, the code will be freely available on GitHub].

#### 2.2.1 Eye Blink Changes

Original ADJUST function classified blinks by using two spatial measures (spatial average difference and spatial variance difference) and a temporal measure (kurtosis) of the components. The spatial average difference was calculated by subtracting the average activity at posterior sites from the average activity at anterior sites. Similarly, the spatial variance difference was calculated by subtracting the variance of activity at posterior sites from the variance of activity at anterior sites. Kurtosis was calculated within each epoch and then averaged across epochs to get a single kurtosis value for each component. In order to improve algorithm performance on geodesic nets, the spatial functions were changed to better classify blink artifacts without introducing too many false alarms. The temporal measure was removed because it did not catch any blinks above and beyond the altered spatial functions.

The new spatial layout functions used the “zscore” function in MATLAB to standardize component activity at each electrode within a given component. Next, the average z-score was computed for five different electrode clusters (Figure 1): left eye region, right eye region, center eye region, central region, and posterior region. Assignment of electrodes to specific clusters were determined based on their location in polar coordinates (radius, theta). As shown in Figure 1, to be in any of the eye clusters, the radius had to be between 0.45 and 0.60. The left eye cluster required a theta value between -60 and 0 degrees whereas the right eye cluster required a theta value between 0 and 60 degrees. The center eye cluster was restricted to theta values between -20 and 20 degrees. To be in the central cluster, the radius had to be less than 0.45 and the absolute value for theta had to be between 35 and 109 degrees. The posterior cluster required a radius less than 0.55 and an absolute theta value greater than 109 degrees.

**Figure 1.**
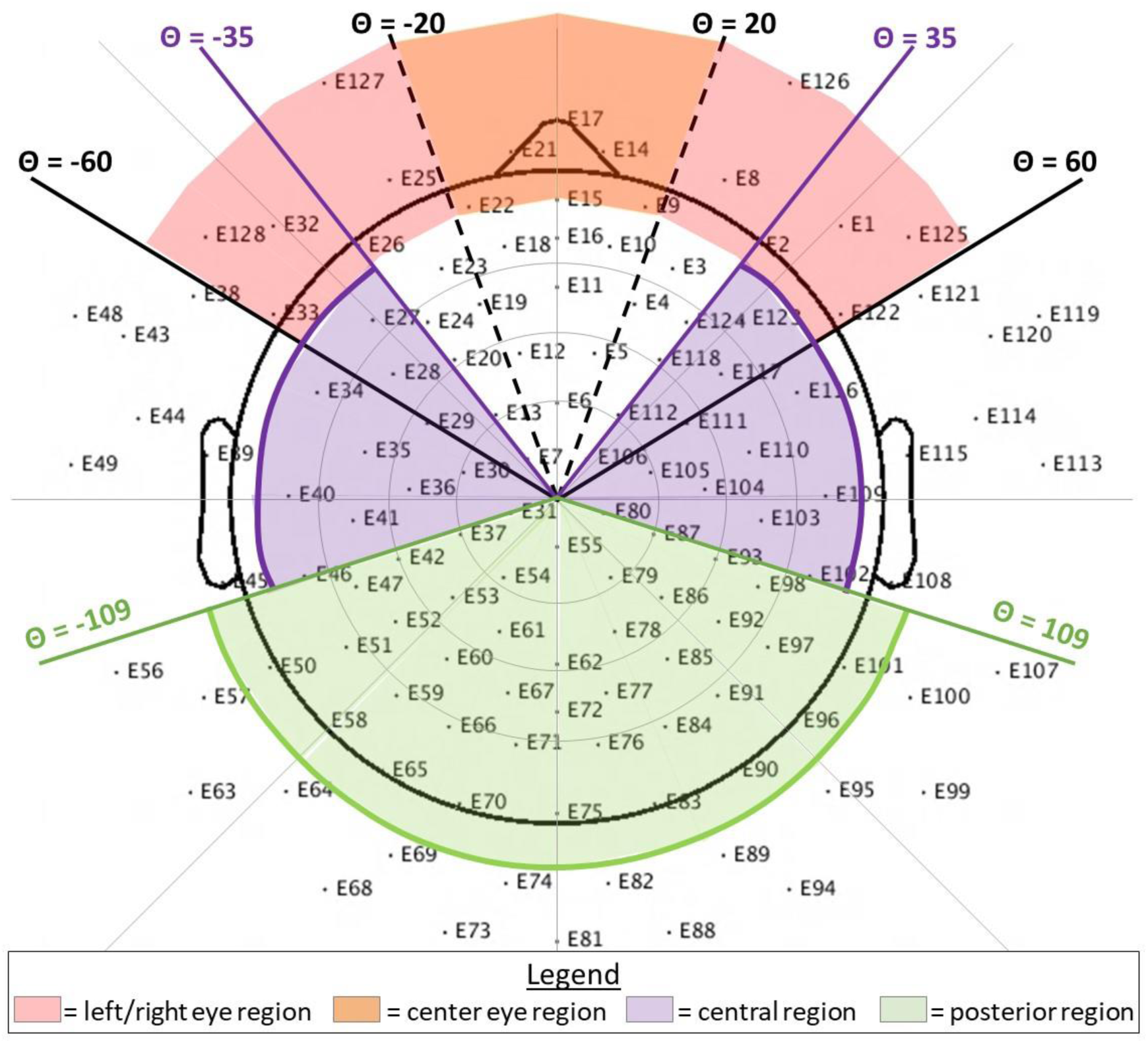
A 128-channel geodesic net layout showing the five regions of interest (left eye, center eye, right eye, central, and posterior) for the new blink detection function. Note that, for the horizontal eye movement detection function, the theta range (+/-35 to +/-62) and radius (>0.5) are slightly different.

In order for a component to be marked as a blink artifact it had to meet three conditions:

1. At least one of the three eye regions (left, right, or center) had to have an average z-score with an absolute value greater than two.
2. The absolute value of the average z-score of the central and posterior regions had to be less than one.
3. The variance of the z-scores in the posterior region had to be less than 0.15 and the variance of the z-scores in the left and right eye regions combined OR the variance of the absolute values of the z-scores in the posterior region had to be less than 0.075 and the variance of the z-scores in the left and right eye regions combined.

Because blinks cause such huge deflections in EEG data at frontal sites (i.e., high activity), the average z-score in components reflecting blinks should well exceed two standard deviations from the mean. Ambiguous components that might reflect neural data, on the other hand, are far less likely to exceed two standard deviations from the mean. For similar reasons, the activity in the back of the head had to be within one standard deviation of the mean and have low variance because blinks should have low activity at posterior sites. Having greater activity at posterior sites would likely indicate the component contains meaningful neural activity.

The new blink detection function was further modified, such that after selecting potential blink components via the three conditions outlined above, secondary checks were added to determine whether blinks had been incorrectly identified based on component spatial spread (from either of the three eye regions towards the central region). Any electrodes that were part of the central eye region or part of the outermost ring of channels were removed before considering spread. Because artifacts from blinks propagate back from the eyes, the algorithm was set up to consider whether or not component activity decreased along the rostral-to-caudal axis. As long as z-scores were less than 2.5 for electrodes with a radius between 0.35 and 0.45 and less than two for electrodes with a radius between 0.25 and 0.35, components were not un-marked as blinks.

#### 2.2.2 Horizontal Eye Movement Changes

Original-ADJUST also classified horizontal eye movements by using a temporal measure (maximum epoch variance) and a spatial measure (spatial eye difference). The spatial eye difference was calculated by taking the absolute value of the difference in activity between the left and right eyes. Maximum epoch variance was calculated by dividing the greatest epoch variance by the average variance across all epochs. Similar to the blink detection code, the temporal function was removed and the spatial function was altered, each for the same reason given in the section entitled “eye blink changes”.

This time, average z-scores were computed for four different electrode clusters: left eye region, right eye region, central region, and posterior region. Electrode assignment to clusters was determined based on the same polar coordinate restrictions that were used for blinks (Figure 1). However, to focus on the lateral electrodes, the left and right eye regions were altered slightly by changing the theta range to be from +/-35 to +/-62 and the radius to be greater than 0.5. In order for a component to be marked as a horizontal eye movement artifact it had to meet three conditions:

1. Either the left or the right eye region had to have an average z-score with an absolute value greater than two
2. The absolute value of the average z-score of the central and posterior regions had to be less than one
3. The variance of the z-scores in the posterior region had to be less than 0.15 and the variance across the left and right eye regions combined had to be greater than 2.5 OR the variance of the absolute values of the z-scores in the posterior region had to be less than 0.075 and the variance across the left and right eye regions combined had to be greater than 2.5

As with blinks, horizontal eye movements cause deflections in EEG data at frontal sites (i.e., high activity) compared to posterior sites, so the first two conditions are the same as described above in the section entitled “eye blink changes”. In the third condition, because saccade components should exhibit opposing polarity along the right-left axis near the eyes (Mognon et al., 2011), the variance of the left and right eye regions had to be greater than 2.5 standard deviations in order for a component to be marked as a saccade. Similar to the new blink detection function, secondary checks were added to determine whether saccades had been incorrectly identified based on component spatial spread (from the ocular region towards the central region). These secondary checks, based on component spread, were identical to those described above for blinks.

#### 2.2.3 Alpha Peak Detector

Many components reflecting neural activity contain local maxima within the alpha band, which we refer to here, for simplicity, as an “alpha peak”. Even though neural activity exists in other power bands (i.e., delta, theta, beta, and gamma), high activity levels in those power bands do not tend to manifest in the EEG power spectrum as a clear peak. As a result, finding high levels of neural activity in power bands outside of the alpha band is not practical; therefore, we chose to focus exclusively on the alpha band.

The ADJUST algorithm was further modified to check for the presence of alpha peaks within any of the potential ICA artifact components. This “alpha peak detector” loops through all of the components marked as artifacts (to be removed from the data) and de-selects any components containing alpha peaks. Code from the MARA toolbox (Winkler, Haufe, & Tangermann, 2011) was adapted to calculate the power spectrum for components using the spectopo function from EEGLAB. Because the EEG power spectrum tends to follow a 1/f curve, the alpha peak detector uses the standardized residual values of the component’s power spectrum after regressing out the estimated 1/f curve. The detector then looks for any local maxima between five Hz and 15 Hz. A larger range for the alpha band was selected due to the fact that alpha tends to be lower in children as compared to adults (Marshall, Bar-Haim, & Fox, 2002) and there are significant individual differences in alpha peaks (Corcoran, Alday, Schlesewsky, & Bornkessel-Schlesewsky, 2018). The detector de-selects components if the local maxima has a prominence greater than 0.3 μV^2^/Hz and a width greater than 0.9 Hz; these thresholds were selected to match one of the expert coders on an independent dataset. More specifically, components from an independent dataset were visually inspected in order to determine which thresholds correctly identified the majority of alpha peaks without an appreciable change in the false alarm rate.

### 2.3 Evaluating adjusted-ADJUST

Performance of the adjusted-ADJUST algorithm was compared to performance of two other algorithms: the original ADJUST algorithm and ICLabel. Performance was measured in three ways: percent agreement with expert coders, number of trials retained after artifact rejection, and reliability (i.e., internal consistency) of the EEG signal. Because the assumption of normality was violated for most variables and the assumption of homogeneity of variances was violated for most comparisons, rank-based non-parametric tests were conducted using the nparLD package (Noguchi, Gel, Brunner, & Konietschke, 2012) in R (v. 3.5.1; R Core Team, 2015). These tests are a non-parametric alternative to the traditional repeated measures ANOVA and they provide a Wald-Type Statistic (WTS) to examine statistical significance. Significant interactions were further probed with planned contrasts using Wilcoxon signed rank tests. In order to control for the false positive rate, a Benjamini-Hochberg procedure (Benjamini & Hochberg, 1995), with a 0.05 false positive discovery rate, was applied to all follow-up comparisons.

#### 2.3.1 Percent Agreement with Expert Coders

As in Mognon and colleagues (2011), three coders with expertise in ICA components derived from EEG classified 3084 components as artifactual or neural. In order for a component to be classified as an artifact, two of the three coders had to classify it as an artifact. Percent agreement scores were then calculated by comparing the artifact selections from each algorithm to the selections of the three expert coders. Additionally, following Mognon et al. (2011), the percent agreement scores were weighted based on how much variance was accounted for by each component. Thus, the percent agreement score (PA) was calculated by dividing the variance accounted for by the components that the algorithm and expert coders agreed on (ICs_Agree_) by the total variance (TV).

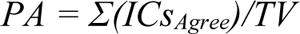

The variance accounted for by each component was calculated using the “pvaf” function in Matlab. This weighted percent agreement score was used because it is more important for the algorithm to correctly classify components that account for the greatest amount of variance in the data. Next, a 3 (Algorithm) x 2 (Age Group) non-parametric test was conducted for the adult and child data using the percent agreement scores. For the infant data, a three-way repeated measures non-parametric test comparing the different algorithms was conducted using the percent agreement scores. Follow-up pairwise comparisons were conducted using a Wilcoxon signed rank test. All post-hoc comparisons were FDR corrected (Benjamini & Hochberg, 1995).

#### 2.3.2 Trials Retained After Removing Artifacts

In order to determine which algorithm not only better matched experts, but also retained more artifact-free data, we compared the results of preprocessing using the adjusted-ADJUST algorithm to the original-ADJUST algorithm, ICLabel, and a preprocessing stream that employed no ICA correction prior to rejecting epochs with ocular activity or excessive noise. This last comparison evaluates the benefits (or disadvantages) of using ICA correction in preprocessing EEG data given that many studies do not employ ICA for artifact correction. The maximum number of epochs that participants could have was 120 (based on the total number of task trials) for dataset 1 and 180 (based on the total number of one-second segments) for dataset 2. After running each algorithm, or not running an algorithm in the case of the no ICA correction condition, the data went through a final artifact rejection step as detailed in the EEG preprocessing section. The proportion of trials remaining after artifact rejection for each condition for children and adults was then used in a 4 (Algorithm) x 2 (Age Group) repeated measures non-parametric test. For infant data, a three-way repeated measures non-parametric test comparing the different conditions was conducted using the proportion of trials remaining after artifact rejection. Follow-up pairwise comparisons were conducted using a Wilcoxon signed rank test. All post-hoc comparisons were FDR corrected (Benjamini & Hochberg, 1995).

#### 2.3.3 Internal Consistency Reliability

Generally, retaining more trials benefits ERP or time frequency analyses by increasing the signal to noise ratio. However, if the trials retained after ICA correction still contain a significant amount of noise, they may decrease the overall data quality rather than increase it. To ensure that including more trials did not reduce the quality of data, the internal consistency reliability was calculated after data correction with adjusted-ADJUST, original-ADJUST, ICLabel, or no ICA correction condition. For adult and child data, we estimated the internal consistency reliability of the mean amplitude of the P1 visual ERP; while, for infant data, we calculated the internal consistency reliability of relative alpha power. The internal consistency reliability estimates were measured via the Spearman-Brown split-half correlation method. Because different methods are expected to differ in the number of artifact-free trials and reliability estimates differ as a function of recording length (i.e., numbers of trials), reliability estimates were obtained for increasing number of trials. This method allowed us to compare the different algorithms while maintaining a constant trial number. Moreover, it allowed us to determine the amount of data necessary to obtain acceptable and excellent reliability. The artifact correction/rejection method that maximizes the signal while reducing noise should be more internally consistent and yield higher reliability estimates even with relatively low amounts of data.

Specifically, internal consistency reliability estimates were obtained using Spearman–Brown-corrected split-half correlation coefficients for an increasing number of trials. From two through nine trials, these correlation coefficients were calculated in steps of one trial; from 10 through 120 trials correlation coefficients were calculated in steps of five trials (i.e., 1, 2, 3, 4, 5, 6, 7, 8, 9, 10, 15, 20, 25, etc.). Following the method proposed by Towers and Allen (2009), for each number of trials, *n*, 10,000 iterations of split-half correlations were calculated. For each iteration, *n* trials were randomly selected from all available trials for that participant. The selected trials were then halved by randomly assigning trials/epochs to one of two groups/halves. The mean P1 amplitude or relative alpha power values were then calculated separately for each half. The Pearson correlation coefficient was obtained using each half across all participants with enough usable trials for any given *n*. Finally, the Spearman-Brown prophecy formula was applied to correct the reliability estimate for test length (i.e. estimate the reliability when the number of trials is doubled given that reliability estimate was calculated with half the original number of trials). This process generated 10,000 reliability estimates for a given *n* for each condition. Notably, when iterating across number of trials (from two to 120), the number of participants included in the split-half reliability estimates decreased as the number of trials increased. Only reliability estimates with a minimum of six participants (the majority of each group) are presented.

For the reliability analyses, whether a method achieved good and excellent reliability for the average reliability coefficients was quantified. Moreover, to provide a measure of the resampling distributions, we present the percentage of iterations that meet or exceed good (.80) and excellent (.90) reliability for each condition. Finally, to quantify differences between conditions, the area under the curve was calculated for each iteration. The average and 95% confidence intervals (CIs) from the resampling distribution were estimated for each condition, providing a measure of overall reliability across increasing numbers of trials – with a greater area under the curve indicating higher internal consistency reliability.

## 3. Results

### 3.1 Agreement with Expert Coders

#### 3.1.1 Adult and Child Data

The results of the 3 (Algorithm) x 2 (Age Group) non-parametric test for the percent agreement scores based on all components can be found in Table 1. As expected, the 3 (Algorithm) x 2 (Age Group) non-parametric test showed an interaction between Algorithm and Age Group, *WTS*(2)=6.62, *p*=0.037. Based on *a priori* hypotheses, planned comparisons were conducted and descriptive information can be found in Table 2. These comparisons revealed that, for children, adjusted-ADJUST performed significantly better than both original-ADJUST, *Z*=2.60, *p*=0.009, *q*=0.016, and ICLabel, *Z*=2.80, *p*=0.005, *q*=0.016. Similarly, in adults, adjusted-ADJUST performed better than original-ADJUST, *Z*=2.60, *p*=0.009, *q*=0.016, and ICLabel, *Z*=2.19, *p*=0.028, *q*=0.036. Comparing ICLabel to original-ADJUST revealed no significant differences for children, *Z*=0.46, *p*=0.646, *q*=0.646, or adults, *Z*=1.38, *p*=0.169, *q*=0.190. As summarized in Figure 2, the adjusted-ADJUST algorithm agreed more with expert coders than both original-ADJUST and ICLabel, for both children and adults. However, as demonstrated by the significant interaction and shown in Figure 2, this difference was larger in children compared to adults.

**Figure 2.**
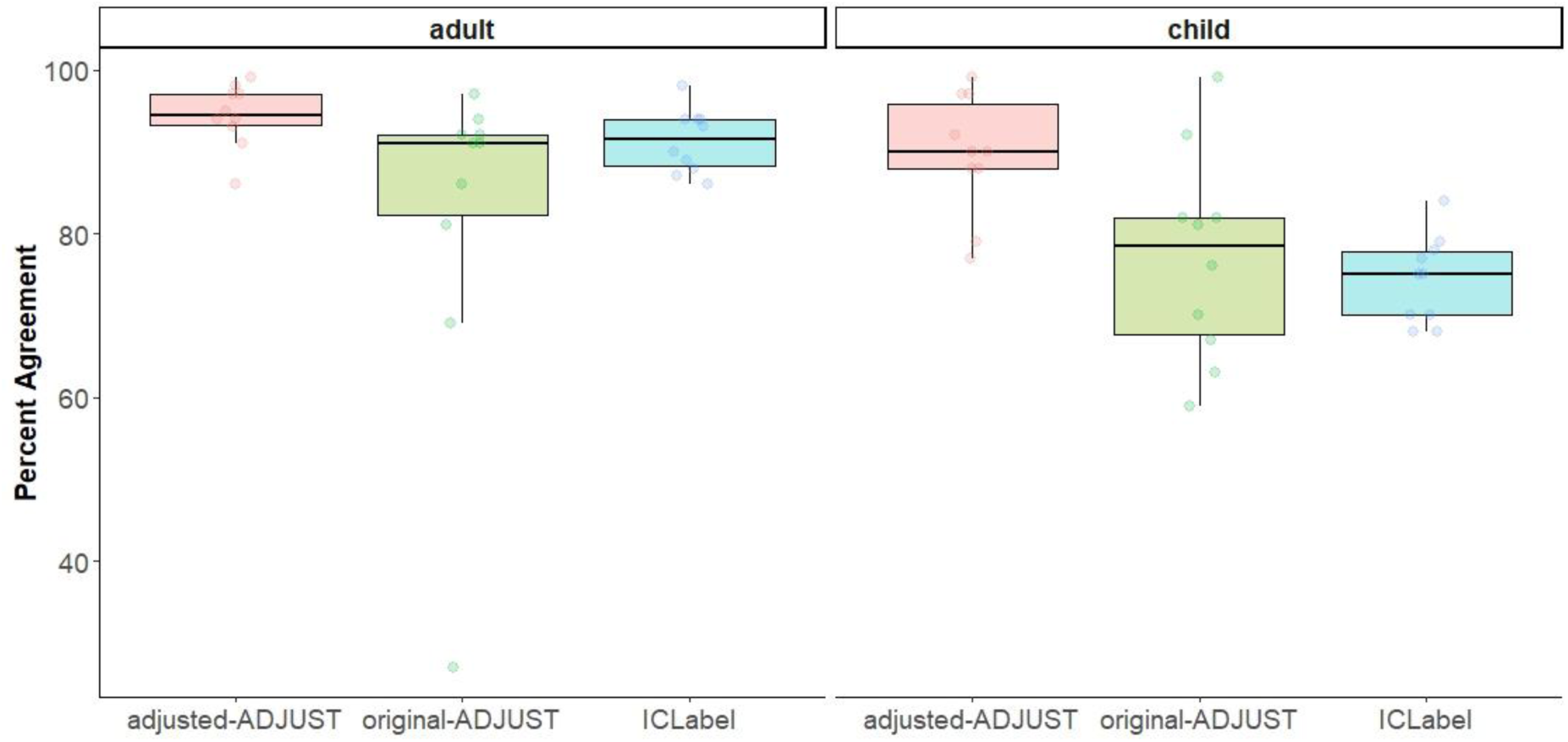
Box plots showing the distribution of the percent agreement scores for children and adults further split up by each algorithm (adjusted-ADJUST, original-ADJUST, or ICLabel). Each dot represents a score.

**Table 1.** Repeated Measures non-parametric test of Percent Agreement Scores in adult and child data.

|  | WTS | df | p-value |
| --- | --- | --- | --- |
| Age | 16.1714 | 1 | 0.0001 |
| Algorithm | 33.1522 | 2 | 0.0000 |
| Age*Algorithm | 6.6191 | 2 | 0.0365 |

**Table 2.** Descriptive information on percent agreement scores between expert coders and the three correction algorithms for adult and child data.

| Age |  | N | Mean | Std. Deviation | Minimum | Maximum | Percentiles |  |  |
| --- | --- | --- | --- | --- | --- | --- | --- | --- | --- |
|  |  |  |  |  |  |  | 25th | 50th (Median) | 75th |
| adult | adjusted-ADJUST | 10 | 94.45% | 3.81% | 86.24% | 98.85% | 92.46% | 94.59% | 97.46% |
|  | original-ADJUST | 10 | 81.99% | 20.86% | 27.48% | 97.08% | 77.67% | 90.97% | 92.81% |
|  | ICLabel | 10 | 91.40% | 3.87% | 85.82% | 98.17% | 88.07% | 91.56% | 94.10% |
| child | adjusted-ADJUST | 10 | 89.57% | 7.36% | 76.53% | 99.35% | 85.69% | 89.80% | 96.52% |
|  | original-ADJUST | 10 | 77.20% | 12.74% | 59.29% | 99.00% | 65.68% | 78.86% | 84.70% |
|  | ICLabel | 10 | 74.47% | 5.35% | 67.64% | 83.98% | 69.52% | 75.16% | 78.57% |

#### 3.1.2 Infant Data

The non-parametric test with infant data showed a marginally significant main effect of Algorithm, *WTS*(2)=5.93, *p*=0.051. Based on *a priori* hypotheses, planned comparisons were conducted and descriptive information can be found in Table 3. These comparisons revealed that adjusted-ADJUST performed significantly better than original-ADJUST, *Z*=2.50, *p*=0.013, *q*=0.039, but not ICLabel, *Z*=- 0.357, *p*=0.721, *q*=0.721. Furthermore, comparing ICLabel to original-ADJUST revealed no significant differences, *Z*=0.866, *p*=0.386, *q*=0.579. As summarized in Figure 3, adjusted-ADJUST agreed more with expert coders than original-ADJUST, but did not differ from ICLabel for the infant data.

**Figure 3.**
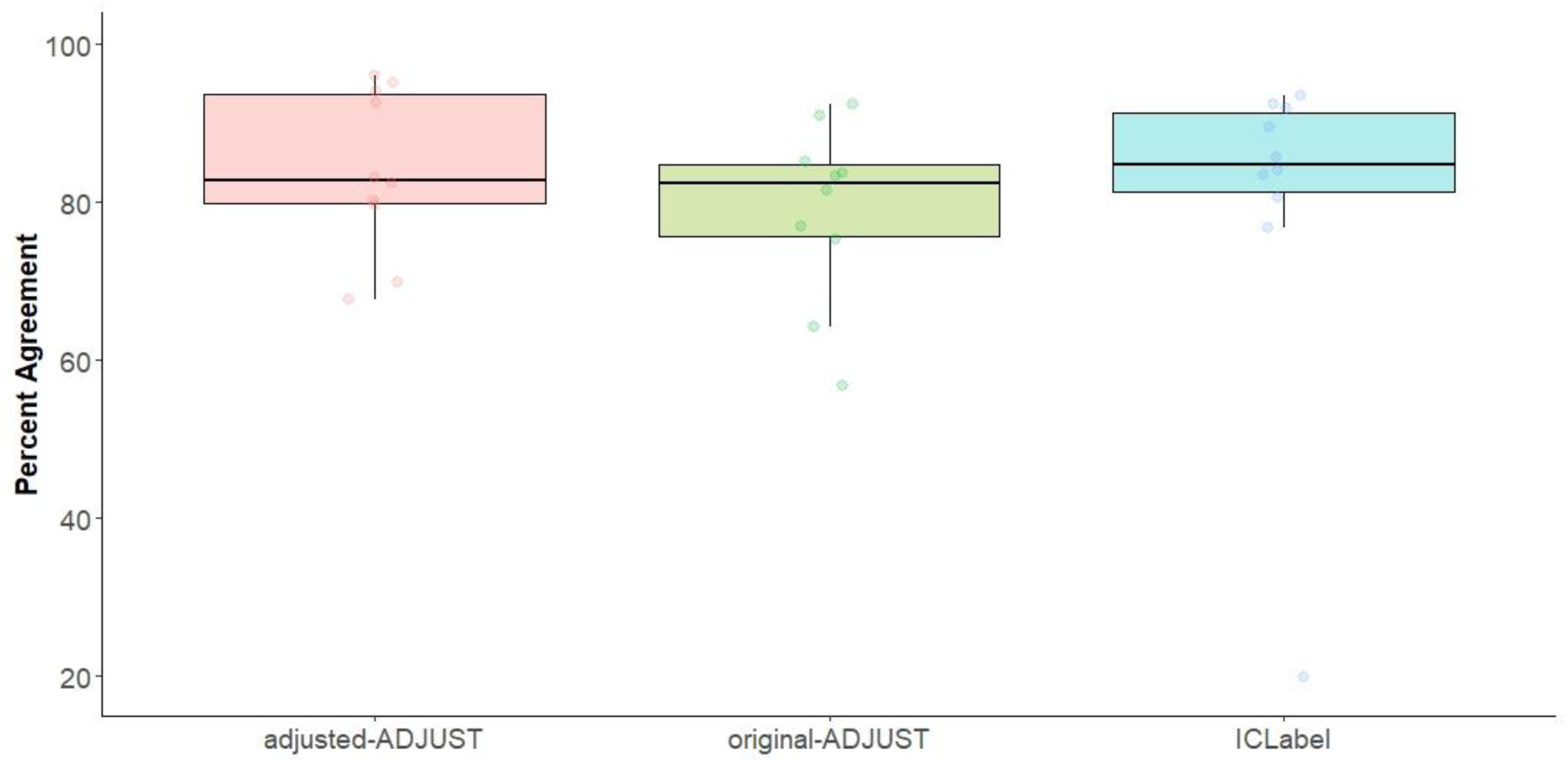
Box plots showing the distribution of the percent agreement scores for infants further split up by each algorithm (adjusted-ADJUST, original-ADJUST, or ICLabel). Each dot represents a score.

**Table 3.** Descriptive information on percent agreement scores between expert coders and the three correction algorithms for infant data.

|  | N | Mean | Std. Deviation | Minimum | Maximum | Percentiles |  |  |
| --- | --- | --- | --- | --- | --- | --- | --- | --- |
|  |  |  |  |  |  | 25th | 50th (Median) | 75th |
| adjusted-ADJUST | 10 | 84.06% | 10.26% | 67.65% | 96.00% | 77.18% | 82.76% | 94.34% |
| original-ADJUST | 10 | 79.01% | 11.22% | 56.83% | 92.44% | 72.58% | 82.39% | 86.54% |
| ICLabel | 10 | 79.75% | 21.73% | 19.89% | 93.52% | 79.56% | 84.80% | 92.00% |

### 3.2 Trials Retained

#### 3.2.1 Adult and Child Data

The results of the 4 (Algorithm) x 2 (Age Group) non-parametric test for trials retained can be found in Table 4 and descriptive information can be found in Table 5. As expected, there was a significant interaction between Algorithm and Age Group, *WTS*(3)=46.74, *p*<0.001, suggesting that the amount of trials retained after using each correction algorithm differed by age group (Figure 4). As a result, planned comparisons were conducted separately for children and adults between adjusted-ADJUST and the other three Algorithm conditions (original-ADJUST, ICLabel, and no ICA correction). These comparisons revealed that adjusted-ADJUST retained significantly more trials than no ICA correction and original- ADJUST for both adults, *Z*=2.80, *p*=0.005, *q*=0.015; *Z*=2.36, *p*=0.018, *q*=0.027, and children, *Z*=2.81, *p*=0.005, *q*=0.015; *Z*=2.67, *p*=0.008, *q*=0.016. There were no significant differences between adjusted-ADJUST and ICLabel for children, *Z*=1.49, *p*=0.137, *q*=0.164, or adults, *Z*=0.412, *p*=0.680, *q*=0.680. These results suggested that, for both children and adults, adjusted-ADJUST retained more trials than both original-ADJUST and no ICA correction, but not ICLabel. However, as shown in Figure 4, the significant interaction suggests that this difference between adjusted-ADJUST and no ICA correction is larger in adults compared to children, while the difference between adjusted-ADJUST and original-ADJUST shows the reverse effect (i.e., the difference is larger in children compared to adults).

**Figure 4.**
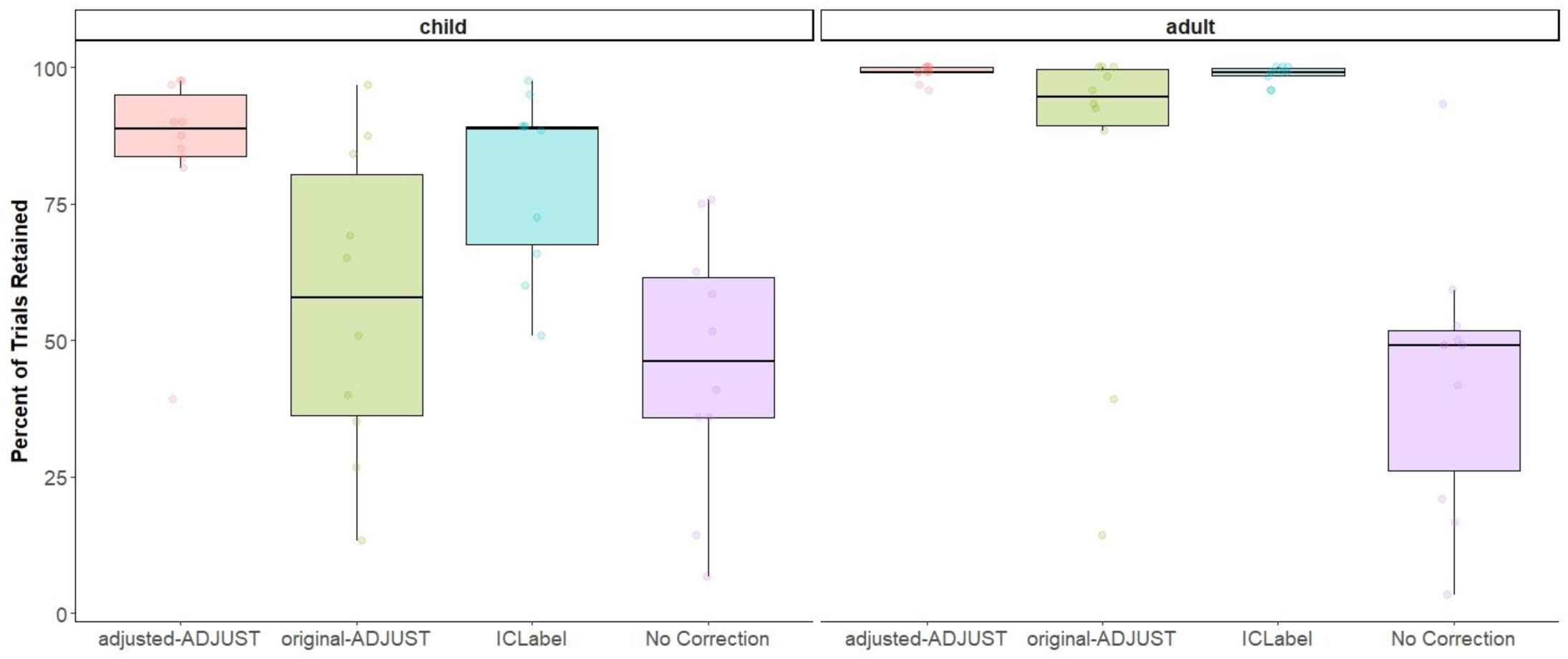
Box plots showing the number of trials retained after artifact rejection for each condition (adjusted-ADJUST, original-ADJUST, ICLabel, or No Correction) further split up by age group (child or adult). Each dot represents a score.

**Table 4.**
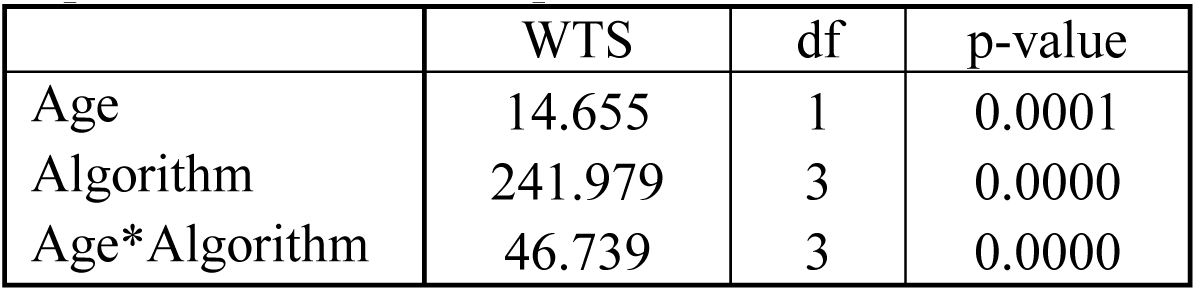
Repeated Measures non-parametric test on the number of trials retained in adult and child data.

**Table 5.**
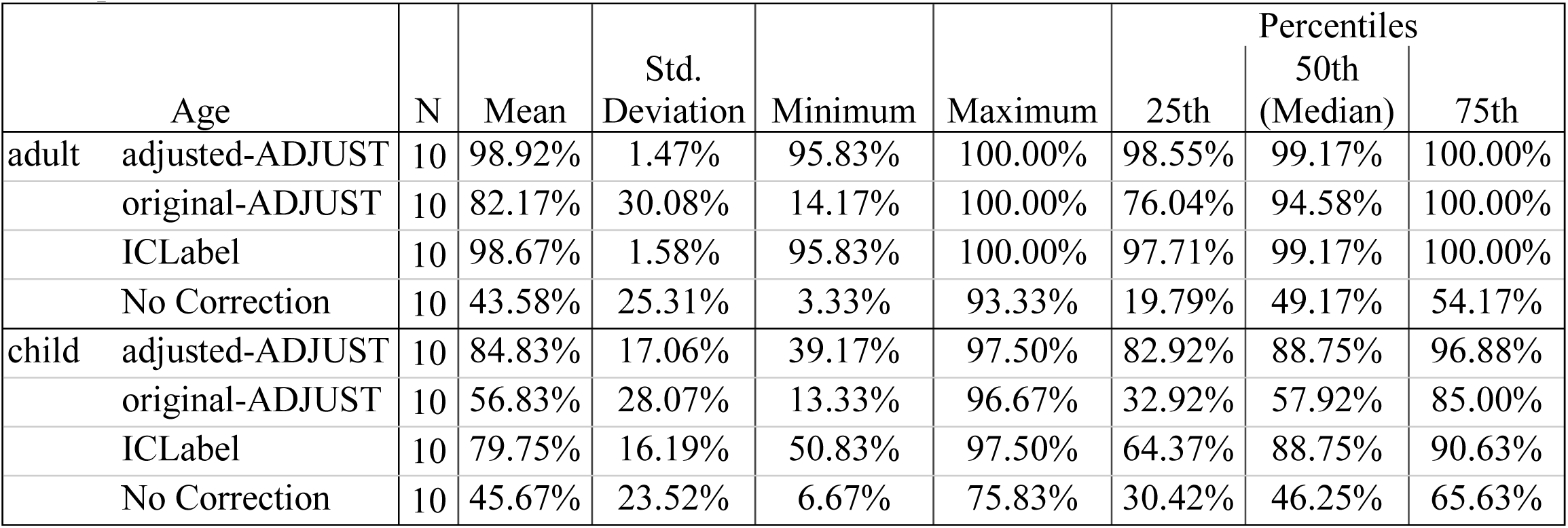
Descriptive information on the number of trials retained with adult and child data

#### 3.2.2 Infant Data

The non-parametric test on trials retained showed a significant main effect of Algorithm, *WTS*(3)=27.16, *p*<0.001. As such, planned comparisons were conducted and descriptive information can be found in Table 6. The adjusted-ADJUST algorithm retained significantly more trials than no ICA correction, *Z*=2.50, *p*=0.013, *q*=0.039. However, there were no significant differences between adjusted-ADJUST and original-ADJUST, *Z*=1.72, *p*=0.086, *q*=0.129, or ICLabel, *Z*=1.27, *p*=0.203, *q*=0.203. These results suggested that, for infants, adjusted-ADJUST retained more trials than no ICA correction, but not original-ADJUST or ICLabel (Figure 5).

**Figure 5.**
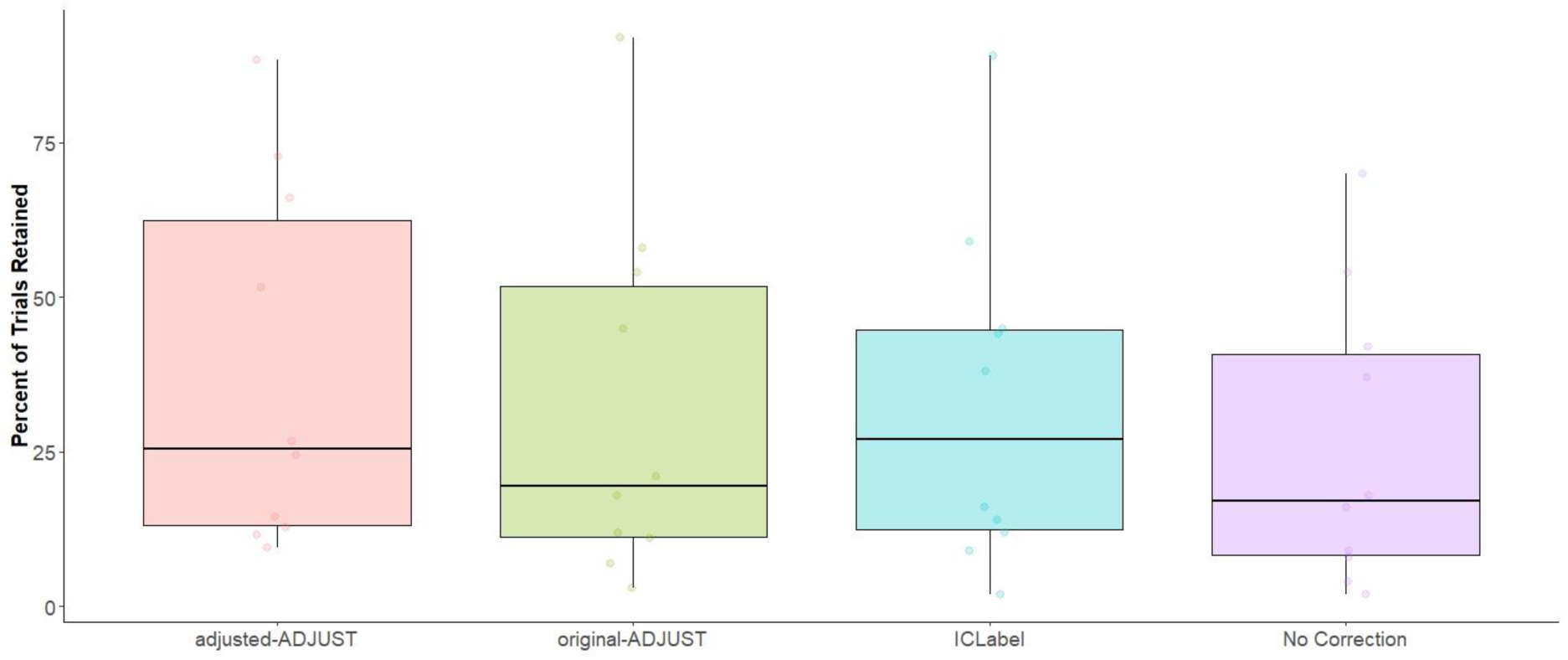
Box plots showing the percent of trials retained after artifact rejection for each condition (adjusted-ADJUST, original-ADJUST, ICLabel, or No Correction) for infants. Each dot represents a score.

**Table 6.** Descriptive information on the number of trials retained with infant data.

|  | N | Mean | Std. Deviation | Minimum | Maximum | Percentiles |  |  |
| --- | --- | --- | --- | --- | --- | --- | --- | --- |
|  |  |  |  |  |  | 25th | 50th (Median) | 75th |
| adjusted-ADJUST | 10 | 37.83% | 29.31% | 9.44% | 88.33% | 12.50% | 25.56% | 67.78% |
| original-ADJUST | 10 | 32.06% | 29.21% | 2.78% | 92.22% | 9.59% | 19.45% | 55.41% |
| ICLabel | 10 | 32.78% | 27.42% | 1.67% | 88.89% | 10.98% | 27.22% | 48.47% |
| No Correction | 10 | 26.11% | 23.46% | 2.22% | 70.00% | 6.95% | 16.95% | 45.28% |

### 3.3 Reliability

Overall, reliability estimates increased with the number of trials. However, this was not always the case. In particular, the no ICA correction condition sometimes exhibited a decline in reliability estimates as the number of trials increased. This decline in reliability is likely the result of significantly less trials being retained (see above), and therefore, less subjects being available for computing reliability estimates at higher trial counts.

For adults, as shown in Figure 6A, all ICA-correction procedures reached a mean of excellent reliability. However, as demonstrated in Figure 6B, when examining the resampling distributions, all ICA-correction procedures reached good reliability (>.8) with a 95% CI, but no method reached excellent (>.9) reliability with a 95% CI. In line with these results, when examining the area under the curve of the average reliability curves (Figure 6C), the 95% CIs of adjusted-ADJUST overlapped with the 95% CIs of the two other ICA-correction methods. However, the CIs for adjusted-ADJUST did not overlap the CIs for no ICA correction, suggesting significantly higher internal consistency reliability.

**Figure 6.**
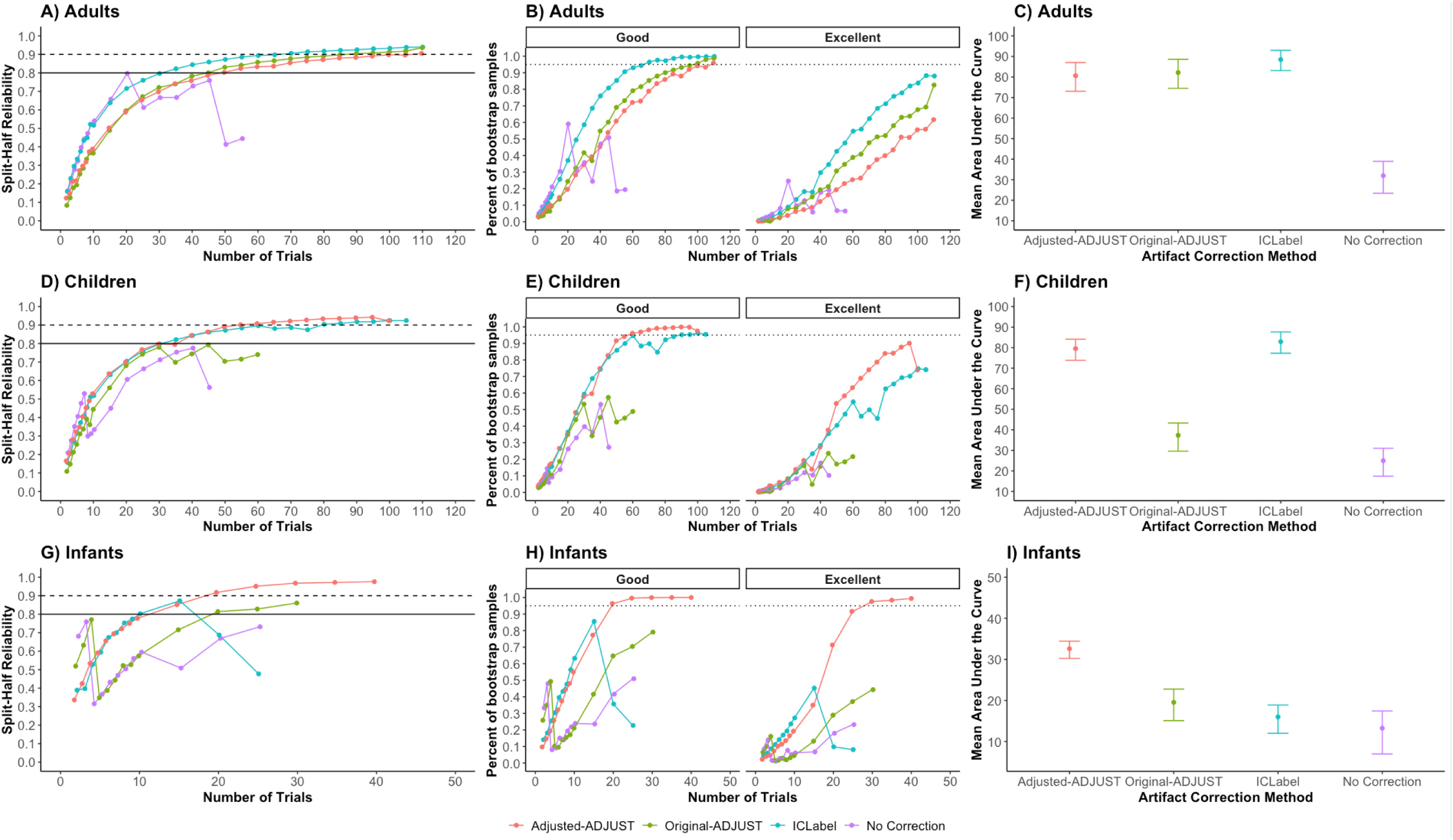
Internal consistency reliability of the mean P1 ERP amplitude in adults and alpha power in infants. Plots A, D, and G show the mean reliability across the 10,000 samples at each number of trials, *n*. Plots B, E, and H show the percentage of the resampling distribution that achieves good (left) and excellent (right) reliability. Plots C, F, and I show the average area under the curve of the reliability curves (plots B, E, and H) with the 95% CIs of the resampling distribution.

For children, as shown in Figure 6D, only adjusted-ADJUST and ICLabel reached an average of excellent reliability, whereas original-ADJUST and no ICA correction only reached good reliability. As demonstrated in Figure 6E, when examining the resampling distributions, adjusted-ADJUST and ICLabel reached good reliability with a 95% CI, while original-ADJUST and no ICA correction did not. Moreover, no method reached excellent reliability with a 95% CI. In line with these results, when examining the area under the curve of the average reliability curves (Figure 6F), the 95% CIs of adjusted-ADJUST overlapped with the 95% CIs of ICLabel. However, the CIs for adjusted-ADJUST did not overlap the CIs for original-ADJUST and no ICA correction, suggesting greater internal consistency reliability.

For infants, as shown in Figure 6G, only adjusted-ADJUST reached a mean of excellent reliability, whereas ICLabel and original-ADJUST reached good reliability, and no ICA correction did not reach good reliability. As demonstrated in Figure 6H, when examining the resampling distributions, only adjusted-ADJUST reached excellent reliability with a 95% CI, while ICLabel, original-ADJUST, and no ICA correction failed to achieve either good or excellent reliability. In line with these results, when examining the area under the curve of the average reliability curves (Figure 6I), the 95% CIs of adjusted-ADJUST did not overlap the 95% CIs of ICLabel, original-ADJUST, and no ICA correction, suggesting greater internal consistency reliability over all other methods tested.

## 4. Discussion

The goal of the present study was to modify an existing artifact selection algorithm to objectively select artifactual components derived from pediatric EEG recorded on geodesic nets. Towards this end, the *original* ADJUST algorithm (Mognon, Jovicich, Bruzzone, & Buiatti, 2011) was optimized to account for the spatial layout of geodesic nets, which are commonly used to collect pediatric EEG, and to de- select components that may include neural activity (e.g., alpha peaks). Our results suggest that the adjusted-ADJUST algorithm performs comparably to or better than original-ADJUST and ICLabel with adult data, but outperformed these existing ICA selection algorithms on several measures when applied to pediatric data. These results suggest that the changes to the ADJUST algorithm for EEG data recorded on geodesic nets improve its performance, especially for pediatric data, and using the adjusted-ADJUST algorithm could facilitate EEG studies with infants and children.

Compared to original-ADJUST, the adjusted-ADJUST algorithm had higher percent agreement scores with expert coders for all age groups and retained more trials for children and adults. Importantly, for children and infants, adjusted-ADJUST also yielded a more reliable signal than original-ADJUST. Furthermore, when compared to a novel algorithm, ICLabel, the adjusted-ADJUST algorithm better matched expert coders on child and adult data. However, across all ages, adjusted-ADJUST did not significantly differ from ICLabel in terms of the number of trials retained. Finally, while adjusted-ADJUST and ICLabel generated similar reliability estimates for children and adults, adjusted-ADJUST yielded significantly greater internal reliably estimates than ICLabel for infants. This difference with infant data is in line with anecdotal evidence, which suggests that ICLabel may have subpar performance in populations not included in the ICLabel training dataset, such as infants (Pion-Tonachini et al., 2019).

Comparing the correction algorithms to the no ICA correction condition suggests that it is better to use some form of ICA correction on EEG data prior to artifact rejection. When assessing trials retained and internal consistency reliability, the correction algorithms generally outperformed the no correction condition by retaining significantly more trials or yielding a higher reliability estimate. This implies that, when trial counts are low, EEG studies would benefit from using ICA-correction methods rather than just deleting trials with artifacts. Moreover, this is not to say that all three correction algorithms are equally beneficial. Performance still varied across the three automated algorithms as some algorithms retained more trials or had greater internal consistency reliability than others.

After considering all three performance measures, it appears that both adjusted-ADJUST and ICLabel could be useful automated algorithms for classifying artifactual components derived from EEG data. However, both algorithms have advantages and limitations that should be considered before implementation. For example, while adjusted-ADJUST and ICLabel performed similarly for adults in terms of trials retained and reliability achieved, adjusted-ADJUST outperformed ICLabel in terms of reliability achieved when applied to infant data. Thus, the adjusted-ADJUST algorithm may currently represent a better method for identifying ICA artifacts in pediatric data. Relatedly, ICLabel uses machine learning to classify components as artifactual or neural, but the training data set that ICLabel uses contains mostly adult data (Pion-Tonachini, Kreutz-Delgado, & Makeig, 2019). To optimize ICLabel for pediatric data, a greater proportion of pediatric data may need to be included in its training dataset. It is also worth noting that some users may prefer an approach that does not use machine learning, as the former approach can allow users to see how data is being classified as artifactual or neural. Moreover, the specific parameters employed in the adjusted-ADJUST algorithm can be modified by users as needed, to be more or less conservative, depending on the population and the preferences of the researchers. For example, if a researcher finds that adjusted-ADJUST is too conservative on eye-blink detection, the z-score value for the frontal eye regions can be modified to match the desired level of blink-identification. Similarly, the thresholds for identifying potential neural activity within a component (i.e., the presence of alpha-peaks) can be adjusted to meet the preference of the researcher. Specifically, thresholds corresponding to the prominence and width of a potential alpha peak, as well as the range of frequencies that are inspected for potential alpha peaks, can all be modified by the user. In summary, if researchers are studying pediatric populations, want to know exactly how their ICA data is being classified as artifactual, or modify their artifact detection thresholds, the adjusted-ADJUST algorithm may currently provide a better solution for users.

One limitation of the current study is that the expert coders used in this study were from the same lab where the algorithm was developed. Consequently, the expert coders are likely biased in favor of the criteria used for the adjusted-ADJUST algorithm (e.g., the default thresholds used to detect alpha peaks) which could have influenced the measure of agreement of with expert coders. For this reason, the trials retained and reliability analyses were included to provide more objective assessments of algorithm performance. Another limitation is that, although expert coders classified over 3000 components, the sample consisted of only 10 adults, 10 children, and 10 infants, which likely resulted in some of the analyses being underpowered. Future work could test the adjusted-ADJUST algorithm in larger samples.

In conclusion, the adjusted-ADJUST algorithm performs similarly to the classification of components by expert coders and retains more trials without compromising reliability. Both pediatric and adult data showed higher percent agreement scores with the adjusted-ADJUST algorithm, as compared to original-ADJUST, while children and adults also showed higher percent agreement scores with adjusted-ADJUST as compared to ICLabel. Furthermore, adjusted-ADJUST retained significantly more trials than no ICA correction for both the pediatric and adult data in addition to retaining more trials than original- ADJUST for both children and adults. The reliability analyses indicated comparable performance across algorithms for adults, better performance for adjusted-ADJUST than original-ADJUST for children and infants, and better performance for adjusted-ADJUST than ICLabel with infants. Thus, the adjusted-ADJUST algorithm is a valuable tool that can assist in the analysis of pediatric EEG data.

## Supporting information

adjusted-ADJUST code

## Conflict of interest statement

The authors have no conflict of interest.

### Acknowledgements

We thank the many research assistants involved in data collection and the participating families without whom the study would not have been possible.

