## Supplementary material for "Adjusting ADJUST: Optimizing the ADJUST Algorithm for Pediatric Data Using Geodesic Nets": adjusted-ADJUST code

% The code below contains 4 different matlab functions that have been appended into one file
% when each function appears in the script, a comment is inserted below that line to show the start of the code from the function

% e.g., "% The functionname(...) is copy and pasted below for reference"

% to indicate the end of the inserted function code, a comment is inserted along with a long line of percent signs (%)

% e.g., "% end of functionname(...)"
         %%%%%%%%%%%%%%%%%%%%%%%%%%%%%%%%

% The final version of the code will be available on GitHub after publication of the manuscript


% adjusted_ADJUST() - Automatic EEG artifact Detector 
% optimized for pediatric populations
%
% Usage:
%   >> [art, horiz, vert, blink, disc,...
%         soglia_DV, diff_var, soglia_K, med2_K, meanK, soglia_SED, med2_SED, SED, soglia_SAD, med2_SAD, SAD, ...
%         soglia_GDSF, med2_GDSF, GDSF, soglia_V, med2_V, nuovaV, soglia_D, maxdin]=adjusted_ADJUST (EEG,out)
% 
% Inputs:
%   EEG        - current dataset structure or structure array (has to be epoched)
%   out        - (string) report file name 
%
% Outputs:
%   art        - List of artifacted ICs
%   horiz      - List of HEM ICs 
%   vert       - List of VEM ICs   
%   blink      - List of EB ICs     
%   disc       - List of GD ICs     
%   soglia_DV  - SVD threshold      
%   diff_var   - SVD feature values
%   soglia_K   - TK threshold      
%   meanK      - TK feature values
%   soglia_SED - SED threshold      
%   SED        - SED feature values
%   soglia_SAD - SAD threshold      
%   SAD        - SAD feature values
%   soglia_GDSF- GDSF threshold      
%   GDSF       - GDSF feature values
%   soglia_V   - MEV threshold      
%   nuovaV     - MEV feature values
%
%

%%%%%%%%%%%%%%%%%%%%%%%%%%%%%%%%%%%%%%%%%%%%%%%%%%%%%%%%%%%%%%%%%%%%%%%%%%%
%
% ADJUST
% Automatic EEG artifact Detector based on the Joint Use of Spatial and Temporal features
% 
% Developed 2007-2014
% Andrea Mognon (1) and Marco Buiatti (2), 
% (1) Center for Mind/Brain Sciences, University of Trento, Italy
% (2) INSERM U992 - Cognitive Neuroimaging Unit, Gif sur Yvette, France
% 
% Last update: 02/05/2014 by Marco Buiatti
%
%%%%%%%%%%%%%%%%%%%%%%%%%%%%%%%%%%%%%%%%%%%%%%%%%%%%%%%%%%%%%%%%%%%%%%%%%%%
% 
% Reference paper:
% Mognon A, Jovicich J, Bruzzone L, Buiatti M, 
% ADJUST: An Automatic EEG artifact Detector based on the Joint Use of Spatial and Temporal features. 
% Psychophysiology 48 (2), 229-240 (2011).
%
%%%%%%%%%%%%%%%%%%%%%%%%%%%%%%%%%%%%%%%%%%%%%%%%%%%%%%%%%%%%%%%%%%%%%%%%%%%
%
% adjusted_ADJUST
% a modified version of ADJUST that has been optimized with pediatric data and geodesic sensor nets
% 
%%%%%%%%%%%%%%%%%%%%%%%%%%%%%%%%%%%%%%%%%%%%%%%%%%%%%%%%%%%%%%%%%%%%%%%%%%%
%
% Reference paper:
% Leach, S.C., Morales, S., Bowers, M. E., Buzzell, G. A., Debnath, R., Beall, D., Fox, N. A., (submitted). Adjusting ADJUST: Optimizing the ADJUST Algorithm for Pediatric Data Using Geodesic Nets. Psychophysiology.
% 
%%%%%%%%%%%%%%%%%%%%%%%%%%%%%%%%%%%%%%%%%%%%%%%%%%%%%%%%%%%%%%%%%%%%%%%%%%%

% Copyright (C) 2009-2014 Andrea Mognon (1) and Marco Buiatti (2), 
% (1) Center for Mind/Brain Sciences, University of Trento, Italy
% (2) INSERM U992 - Cognitive Neuroimaging Unit, Gif sur Yvette, France
%
% This program is free software; you can redistribute it and/or modify
% it under the terms of the GNU General Public License as published by
% the Free Software Foundation; either version 2 of the License, or
% (at your option) any later version.
%
% This program is distributed in the hope that it will be useful,
% but WITHOUT ANY WARRANTY; without even the implied warranty of
% MERCHANTABILITY or FITNESS FOR A PARTICULAR PURPOSE.  See the
% GNU General Public License for more details.
%
% You should have received a copy of the GNU General Public License
% along with this program; if not, write to the Free Software
% Foundation, Inc., 59 Temple Place, Suite 330, Boston, MA  02111-1307  USA

%%%%%%%%%%%%%%%%%%%%%%%%%%%%%%%%%%%%%%%%%%%%%%%%%%%%%%%%%%%%%%%%%%%%%%%%%%%

% VERSIONS LOG
%
% 02/05/14: Modified text in Report.txt (MB).
%
% 30/03/14: Removed 'message to the user' (redundant). (MB)
% 
% 22/03/14: kurtosis is replaced by kurt for compatibility if signal processing
%           toolbox is missing (MB).
%
% V2 (07 OCTOBER 2010) - by Andrea Mognon
% Added input 'nchannels' to compute_SAD and compute_SED_NOnorm;
% this is useful to differentiate the number of ICs (n) and the number of
% sensors (nchannels);
% bug reported by Guido Hesselman on October, 1 2010.

%%%%%%%%%%%%%%%%%%%%%%%%%%%%%%%%%%%%%%%%%%%%%%%%%%%%%%%%%%%%%%%%%%%%%%%%%%%


% function [art, horiz, vert, blink, disc,...
%         soglia_DV, diff_var, soglia_K, meanK, soglia_SED, SED, soglia_SAD, SAD, ...
%         soglia_GDSF, GDSF, soglia_V, nuovaV, soglia_D, maxdin]=ADJUST (EEG,out)
function [art, horiz, vert, blink, disc,...
        soglia_DV, diff_var, soglia_K, med2_K, meanK, soglia_SED, med2_SED, SED, soglia_SAD, med2_SAD, SAD, ...
        soglia_GDSF, med2_GDSF, GDSF, soglia_V, med2_V, nuovaV, soglia_D, maxdin]=adjusted_ADJUST (EEG,out)

    
%% Settings

% ----------------------------------------------------
% |  Change experimental settings in this section    |
% ----------------------------------------------------

% ----------------------------------------------------
% |  Initial message to user:                        |
% ----------------------------------------------------
% 
% disp(' ')
% disp('Detects Horizontal and Vertical eye movements,')
% disp('Blinks and Discontinuities in dataset:')
% disp([EEG.filename])
% disp(' ')

% ----------------------------------------------------
% |  Collect useful data from EEG structure          |
% ----------------------------------------------------

%number of ICs=size(EEG.icawinv,1);

%number of time points=size(EEG.data,2);

if length(size(EEG.data))==3
    
    num_epoch=size(EEG.data,3);
       
else
    
    num_epoch=0;

end

% Check the presence of ICA activations

if isempty(EEG.icaact)
    disp('EEG.icaact not present. Recomputed from data.');
    if length(size(EEG.data))==3
%         EEG.icaact = EEG.icaweights*EEG.icasphere*reshape(EEG.data, size(EEG.icawinv,1), num_epoch*size(EEG.data,2));
%         EEG.icaact = reshape(EEG.icaact,size(EEG.icawinv,1),size(EEG.data,2), num_epoch);
         EEG.icaact = reshape(EEG.icaweights*EEG.icasphere*reshape(EEG.data,[size(EEG.data,1)...
 size(EEG.data,2)*size(EEG.data,3)]),[size(EEG.data,1) size(EEG.data,2) size(EEG.data,3)]);
    else EEG.icaact = EEG.icaweights*EEG.icasphere*EEG.data;
    end
end

topografie=EEG.icawinv'; %computes IC topographies

% Topographies and time courses normalization
% 
% disp(' ');
% disp('Normalizing topographies...')
% disp('Scaling time courses...')

for i=1:size(EEG.icawinv,2) % number of ICs
    
    ScalingFactor=norm(topografie(i,:));
    
    topografie(i,:)=topografie(i,:)/ScalingFactor;
 
    if length(size(EEG.data))==3
        EEG.icaact(i,:,:)=ScalingFactor*EEG.icaact(i,:,:);
    else
        EEG.icaact(i,:)=ScalingFactor*EEG.icaact(i,:);
    end
    
end
% 
% disp('Done.')
% disp(' ')

% Variables memorizing artifacted ICs indexes

blink=[];

horiz=[];

vert=[];

disc=[];

%% Check EEG channel position information
nopos_channels=[];
for el=1:length(EEG.chanlocs)
    if(any(isempty(EEG.chanlocs(1,el).X)&isempty(EEG.chanlocs(1,el).Y)&isempty(EEG.chanlocs(1,el).Z)&isempty(EEG.chanlocs(1,el).theta)&isempty(EEG.chanlocs(1,el).radius)))
        nopos_channels=[nopos_channels el];
    end;
end

if ~isempty(nopos_channels)
    warning(['Channels ' num2str(nopos_channels) ' have incomplete location information. They will NOT be used to compute ADJUST spatial features']);
    disp(' ');
end;

pos_channels=setdiff(1:length(EEG.chanlocs),nopos_channels);

%% Feature extraction

disp(' ')
disp('Features Extraction:')

%GDSF - General Discontinuity Spatial Feature

disp('GDSF - General Discontinuity Spatial Feature...')

GDSF = compute_GD_feat(topografie,EEG.chanlocs(1,pos_channels),size(EEG.icawinv,2));


%SED - Spatial Eye Difference

disp('SED - Spatial Eye Difference...')

[SED,medie_left,medie_right]=computeSED_NOnorm(topografie,EEG.chanlocs(1,pos_channels),size(EEG.icawinv,2)); 


%SAD - Spatial Average Difference

disp('SAD - Spatial Average Difference...')

[SAD,var_front,var_back,mean_front,mean_back]=computeSAD(topografie,EEG.chanlocs(1,pos_channels),size(EEG.icawinv,2));


%SVD - Spatial Variance Difference between front zone and back zone

diff_var=var_front-var_back;

%epoch dynamic range, variance and kurtosis

K=zeros(num_epoch,size(EEG.icawinv,2)); %kurtosis
Kloc=K;

Vmax=zeros(num_epoch,size(EEG.icawinv,2)); %variance

% disp('Computing variance and kurtosis of all epochs...')

for i=1:size(EEG.icawinv,2) % number of ICs
    
    for j=1:num_epoch              
        Vmax(j,i)=var(EEG.icaact(i,:,j));        
%         Kloc(j,i)=kurtosis(EEG.icaact(i,:,j));
        K(j,i)=kurt(EEG.icaact(i,:,j));
    end  
end

% check that kurt and kurtosis give the same values:
% [a,b]=max(abs(Kloc(:)-K(:)))

%TK - Temporal Kurtosis

disp('Temporal Kurtosis...')

meanK=zeros(1,size(EEG.icawinv,2));

for i=1:size(EEG.icawinv,2)
    if num_epoch>100
    meanK(1,i)=trim_and_mean(K(:,i)); 
    else meanK(1,i)=mean(K(:,i));
    end

end


%MEV - Maximum Epoch Variance

disp('Maximum epoch variance...')

maxvar=zeros(1,size(EEG.icawinv,2));
meanvar=zeros(1,size(EEG.icawinv,2));


for i=1:size(EEG.icawinv,2)
    if num_epoch>100
     maxvar(1,i)=trim_and_max(Vmax(:,i)');
     meanvar(1,i)=trim_and_mean(Vmax(:,i)');
    else 
     maxvar(1,i)=max(Vmax(:,i));
     meanvar(1,i)=mean(Vmax(:,i));
    end
end

% MEV in reviewed formulation:

nuovaV=maxvar./meanvar;


%% Thresholds computation

disp('Computing EM thresholds...')

% soglia_K=EM(meanK);
% 
% soglia_SED=EM(SED);
% 
% soglia_SAD=EM(SAD);
% 
% soglia_GDSF=EM(GDSF);
% 
% soglia_V=EM(nuovaV); 
[soglia_K,med1_K,med2_K]=EM(meanK);

[soglia_SED,med1_SED,med2_SED]=EM(SED);

[soglia_SAD,med1_SAD,med2_SAD]=EM(SAD);

[soglia_GDSF,med1_GDSF,med2_GDSF]=EM(GDSF);

[soglia_V,med1_V,med2_V]=EM(nuovaV); 

%% Output file header

% ----------------------------------------------------
% |  Opens report file and writes header             |
% ----------------------------------------------------

file=fopen(out,'w');

fprintf(file,'adjusted_ADJUST\n');

fprintf(file,'Automatic EEG artifacts Detector optimized for pediatric data\n\n');

fprintf(file,'adapted from Andrea Mognon and Marco Buiatti (2009-2014)\n\n');

fprintf(file,['Analyzed dataset: ' EEG.filename '\n']);

fprintf(file,['Analysis date: ' date '\n']);

fprintf(file,'Analysis carried out on the %d Independent Components\n\n',size(EEG.icawinv,2));

%%
%%%%%%%%%%%%%%%%%%%%%%%%%%%%%%%%%%%%%%%%%%%%%%%%%%%%%%%%%%%%%%%%%%%%%%%%%
% adjusted_ADJUST modification                                        %%%
% find all ICs with bumps btw 5 and 15 Hz                             %%%
cwb=MARA_extract_time_freq_features(EEG);                             %%%
%%%%%%%%%%%%%%%%%%%%%%%%%%%%%%%%%%%%%%%%%%%%%%%%%%%%%%%%%%%%%%%%%%%%%%%%%

% MARA_extract_time_freq_features(EEG) function is copy and pasted below for reference

function [features] = MARA_extract_time_freq_features(EEG)
% function [features, values_m, values_a] = MARA_extract_time_freq_features(EEG)    
%      
%   features = list of components with bumps
%                           
data = EEG.data;
fs = EEG.srate;

% transform epoched data into continous data
if length(size(data)) == 3
    s = size(data); 
    data = reshape(data, [EEG.nbchan, prod(s(2:3))]); 
end

%downsample (to 100-200Hz) 
factor = max(floor(EEG.srate/100),1); 
data = data(:, 1:factor:end); 
fs = round(fs/factor); 

%compute icaactivation and standardise variance to 1
icacomps = (EEG.icaweights * EEG.icasphere * data)';
icacomps = icacomps./repmat(std(icacomps,0,1),length(icacomps(:,1)),1);
icacomps = icacomps';

%creating a matrix to help us check for peaks in data
change=[0 linspace(-1,1,50)];
components_with_bumps = [];
components_with_bumps_2 = [];
components_with_plats = [];
components_with_plats_2 = [];
pxx_matrix = zeros(length(icacomps(:,1)),51);
resids_pxx_matrix = zeros(length(icacomps(:,1)),50);

%run through all the components
for ic=1:length(icacomps(:,1)) 
    fprintf('.');
    %%%%%%%%%%%%%%%%%%%%%%%%%%%%%%%%%%%
    % Proc Spectrum for Channel
    %%%%%%%%%%%%%%%%%%%%%%%%%%%%%%%%%%%
    [pxx,freq,speccomp,contrib,specstd] = spectopo(icacomps(ic,:), fs, fs);
    pxx = 10*log10(pxx * 100/2);
    pxx_matrix(ic,1:51)=pxx(1:51);
    %%%%%%%%%%%%%%%%%%%%%%%%%%%%%%%%%%%
    % Get the residual values
    %%%%%%%%%%%%%%%%%%%%%%%%%%%%%%%%%%%
    pxx = real(pxx); %need for spectopo
    pxx = pxx*-1; %need for spectopo
    OneOverF = (1./freq);
    lm_pxx = fitlm(OneOverF(2:51),pxx(2:51));
    resids_pxx = lm_pxx.Residuals.Standardized;
    resids_pxx_matrix(ic,1:50)=resids_pxx;
    

    %%%%%%%%%%%%%%%%%%%%%%%%%%%%%%%%%%%
    % Checking for bumps between 5 and 15 Hz
    %%%%%%%%%%%%%%%%%%%%%%%%%%%%%%%%%%%
    % look for peaks with the residual values
    [pks, locs, widths, proms] = findpeaks(resids_pxx(4:16),[4:16]); %looks for peaks btw 5 and 15 Hz (1 less and more so any bumps at 5 or 15 are caught)
    for numPeaks = 1:length(pks) %in case we have mult. peaks
        if (proms(numPeaks) > 0.3 && pks(numPeaks)>1 && widths(numPeaks) > 0.9) %conservative
%         if (proms(numPeaks) > 0.5 && pks(numPeaks)>1 && widths(numPeaks) > 0.9) %liberal
            components_with_bumps = [components_with_bumps ic];
        end
    end
    
    %%%%%%%%%%%%%%%%
    % We are not sure if this chunk without the residuals is needed... 
    % the code above just seemed too cautious
    % We are testing it now and might make changes
    
    % look for peaks with the NON-residual values
    % much more liberal threshold
    % above code using the residual values sometimes gets tiny plateaus or noise
    % the below code doesn't catch that stuff
    [pks, locs, widths, proms] = findpeaks(pxx(5:17),[5:17]); %looks for peaks btw 5 and 15 Hz (1 less and more so any bumps at 5 or 15 are caught)
    for numPeaks = 1:length(pks) %in case we have mult. peaks
        if (proms(numPeaks) > 0.15) %conservative
%         if (proms(numPeaks) > 0.35) %liberal
            components_with_bumps_2 = [components_with_bumps_2 ic];
        end
    end
    %%%%%%%%%%%%%%%
    
    %%%%%%%%%%%%%%%%%%%%%%%%%%%%%%%%%%%
    % Checking for plateaus between 5 and 15 Hz for first 7 components
    %%%%%%%%%%%%%%%%%%%%%%%%%%%%%%%%%%%
    if ic < 8
        [pks, locs, widths, proms] = findpeaks(resids_pxx(4:16),[4:16]);
        for numPeaks = 1:length(pks) %in case we have mult. peaks
            if (proms(numPeaks) > 0.15 && pks(numPeaks)>1) %conservative
%             if (proms(numPeaks) > 0.15 && pks(numPeaks)>1) %liberal
                components_with_plats = [components_with_plats ic];
            end
        end
        
        %%%%%%%%%%%%%%%%
        % We are not sure if this chunk without the residuals is needed... 
        % the code above just seemed too cautious
        % We are testing it now and might make changes
        [pks, locs, widths, proms] = findpeaks(pxx(5:17)+(10*change(5:17)),[5:17]);
        for numPeaks = 1:length(pks) %in case we have mult. peaks
            if (proms(numPeaks) > 0.05) %consevative
%             if (proms(numPeaks) > 0.05) %liberal
                components_with_plats_2 = [components_with_plats_2 ic];
            end
        end
        %%%%%%%%%%%%%%%
        
    end
    %%%%%%%%%%%%%%%%%%%%%%%%%%%%%%%%%%%%
    %Append Features 
    %%%%%%%%%%%%%%%%%%%%%%%%%%%%%%%%%%%
    %features(:,ic)= [Hz8_13]; 
end

% intialize these for later
features = []; % this will be our output matrix (our list of components with bumps)
features_2 = [];
% combine the two components with bumps matrices
components_with_bumps = unique(components_with_bumps);
components_with_bumps_2 = unique(components_with_bumps_2);
for check=1:length(components_with_bumps)
    if sum(components_with_bumps(check) == components_with_bumps_2)
        % if they both have a component, add it to the features matrix
        features = [features components_with_bumps(check)];
    end
end
% combine the two components with plateaus matrices
components_with_plats = unique(components_with_plats);
components_with_plats_2 = unique(components_with_plats_2);
for check=1:length(components_with_plats)
    if sum(components_with_plats(check) == components_with_plats_2)
        features_2 = [features_2 components_with_plats(check)];
    end
end
% combine the components with bumps matrix and the plateau matrix
features = unique([features features_2]); %output matrix (list of components with bumps)

% plot componetns with bumps and components w/o bumps
figure; hold on;
bumps=subplot(2,1,1);
for jj=1:length(features)
    plot(bumps,1:50,resids_pxx_matrix(features(jj),:))
    hold on
end
title(bumps,'ICs with Bumps')

good_ics=1:EEG.nbchan;
good_ics(features)=[];
nobumps=subplot(2,1,2);
for gg=1:length(good_ics)
    plot(nobumps,1:50,resids_pxx_matrix(good_ics(gg),:))
    hold on
end
title(nobumps,'ICs without Bumps')

par_id=strsplit(EEG.setname,'_')
saveas(gcf,[par_id{1} '.jpg'])

%open a new figure for the spectopo output
hold off; figure;

% features = unique(components_with_bumps);
disp('.');
end

% end of MARA_extract_time_freq_features(EEG) function
%%%%%%%%%%%%%%%%%%%%%%%%%%%%%%%%%%%%%%%%%%%%%%%%%%%%%%%%%%%%%%%%%%%%%%%%%

%% Horizontal eye movements (HEM)

disp(' ');
disp('Artifact Identification:');
disp('Horizontal Eye Movements...')

% ----------------------------------------------------
% |  Writes HEM header in the report file            |
% ----------------------------------------------------

fprintf(file,'> HEM - Horizontal movements\n\n');

fprintf(file,'Classification based on features:\n');

fprintf(file,'SED - Spatial eye difference (threshold=%f)\n',soglia_SED);

fprintf(file,'MEV - Maximum epoch variance (threshold=%f)\n\n',soglia_V);

fprintf(file,'ICs with Horizontal eye movements:\n');

%%%%%%%%%%%%%%%%%%%%%%%%%%%%%%%%%%%%%%%%%%%%%%%%%%%%%%%%%%%%%%%%%%%%%%%%%
% adjusted_ADJUST modification                                        %%%
% horizontal eye movement detection code                              %%%
horiz=beall_horizontal(cwb,topografie,EEG.chanlocs(1,pos_channels),size(EEG.icawinv,2));

% beall_horizontal function is copy and pasted below for reference

% Author: Daniel J. Beall
%
%
% beall_horizontal() - Returns list of ICA components marked with
%                      horizontal eye movement
%
% Usage:
%   >> [horizontal] = beall_horizontal(cwb,topog,chanlocs,n);
%
% Inputs:
%   cwb        - list of components with bumps
%   topog      - topographies vector
%   chanlocs   - EEG.chanlocs struct
%   n          - number of ICs
%   nchannels  - number of channels
% Outputs:
%   horizontal - list of detected blinks
%

function [horizontal] = beall_horizontal(cwb,topog,chanlocs,n)

horizontal = [];
nchannels=length(chanlocs);

%% Get Z-Scores Matrix
zmatrix = zeros(n,n);
for i=1:n % for each topography 
    curr_vect = zeros(n);
    %deep copy ICA matrix
    for h=1:n
        zmatrix(i,h) = topog(i,h);
    end
    for h=1:n
        curr_vect(h,1) = topog(i,h);
    end
    z_vect = zscore(curr_vect);
    for h=1:n
        zmatrix(i,h) = z_vect(h,1);
    end    
end
%            _                       _
% zmatrix = | z11    z12    z13   ... | row1 = IC1
%           | z21    z22    z23   ... | row2 = IC2
%           | z31    z32    z33   ... | ...
%           |_...    ...    ...   zij_|
%          col1=E1  col2=E2 ...             nxn matrix (n=#components)

%% Define scalp zones

% Find electrodes in Frontal Area (FA)
dimlefteyes=0; %number of electrodes
index1=zeros(1,nchannels); %indexes of electrodes
for k=1:nchannels
   if (chanlocs(1,k).theta > -62 && chanlocs(1,k).theta < -35) && (chanlocs(1,k).radius>0.5)
        dimlefteyes=dimlefteyes+1; %count electrodes
        index1(1,dimlefteyes)=k;
    end
end

dimrighteyes=0; %number of electrodes
index2=zeros(1,nchannels); %indexes of electrodes
for k=1:nchannels
   if (chanlocs(1,k).theta < 62 && chanlocs(1,k).theta > 35) && (chanlocs(1,k).radius>0.5) %electrodes are in FA
        dimrighteyes=dimrighteyes+1; %count electrodes
        index2(1,dimrighteyes)=k;
    end
end

% Find electrodes in everywhere else
dimcenter=0; %number of electrodes
indexc=zeros(1,nchannels); %indexes of electrodes
for k=1:nchannels
    if( (abs(chanlocs(1,k).theta)) > 35 && abs(chanlocs(1,k).theta) < 109 && (chanlocs(1,k).radius<0.45) )
        dimcenter=dimcenter+1; %count electrodes
        indexc(1,dimcenter)=k;
    end
end

dimbackleft=0; %number of electrodes
indexbl=zeros(1,nchannels); %indexes of electrodes
for k=1:nchannels
   if (chanlocs(1,k).theta <= -109) && (chanlocs(1,k).radius<0.55) %electrodes are in FA
        dimbackleft=dimbackleft+1; %count electrodes
        indexbl(1,dimbackleft)=k;
    end
end

dimbackright=0; %number of electrodes
indexbr=zeros(1,nchannels); %indexes of electrodes
for k=1:nchannels
   if (chanlocs(1,k).theta >= 109) && (chanlocs(1,k).radius<0.55) %electrodes are in FA
        dimbackright=dimbackright+1; %count electrodes
        indexbr(1,dimbackright)=k;
    end
end

%% z-scores of the front and back

% NOTE: 
% ica struct is used for debugging purposed only
% and has no impact on the actual script run,
% just ignore it.

horizontalCount = 1;
for i=1:n % for each topography
    
    % create FA electrodes vector
    zfrontlefteyes=zeros(1,dimlefteyes);
    for h=1:dimlefteyes
        zfrontlefteyes(1,h)=zmatrix(i,index1(1,h));
    end
    ica(i).lefteyeArr = zfrontlefteyes;
    
    zfrontrighteyes=zeros(1,dimrighteyes);
    for h=1:dimrighteyes
        zfrontrighteyes(1,h)=zmatrix(i,index2(1,h));
    end
    ica(i).righteyeArr = zfrontrighteyes;
    
    % create other electrodes vectors
    zcenter=zeros(1,dimcenter);
    for h=1:dimcenter
        zcenter(1,h)=zmatrix(i,indexc(1,h));
    end
    ica(i).chArr = zcenter;
    
    zbackl=zeros(1,dimbackleft);
    for h=1:dimbackleft
        zbackl(1,h)=zmatrix(i,indexbl(1,h));
    end
    ica(i).blArr = zbackl;

    zbackr=zeros(1,dimbackright);
    for h=1:dimbackright
        zbackr(1,h)=zmatrix(i,indexbr(1,h));
    end
    ica(i).brArr = zbackr;
   
    % Calculates the mean of each z-score vector.
    % Sums the absolute value of each element in the vector,
    % then divides that by the number of elements.
    le = 0; 
    for j=1:dimlefteyes
        le = le + abs(zfrontlefteyes(1,j)); 
    end
    le = le/dimlefteyes;
    ica(i).lemean = le;
    
    re = 0; 
    for j=1:dimrighteyes
        re = re + abs(zfrontrighteyes(1,j)); 
    end
    re = re/dimrighteyes;
    ica(i).remean = re;

    ch = 0; 
    for j=1:dimcenter
        ch = ch + abs(zcenter(1,j)); 
    end
    ch = ch/dimcenter;
    ica(i).chmean = ch;

    bl = 0; 
    for j=1:dimbackleft
        bl = bl + abs(zbackl(1,j)); 
    end
    bl = bl/dimbackleft;
    ica(i).blmean = bl;

    br = 0; 
    for j=1:dimbackright
        br = br + abs(zbackr(1,j)); 
    end
    br = br/dimbackright;
    ica(i).brmean = br;

    % if the left or right side pass a certain threshold and all of the 
    % back are below a certain threshold, then we reject
    if(abs(le) > 2 || abs(re) > 2) %if high activity in front
        if (mean([abs(bl) abs(br) abs(ch)]) < 1) %if low activity in back 
            if((var([zbackl zbackr]')<0.15 || var([abs(zbackl) abs(zbackr)]')<0.075) && var([zfrontlefteyes zfrontrighteyes]')>2.5) %if low variance in back and high variance in front
                horizontal(horizontalCount) = i;
                horizontalCount = horizontalCount + 1;
            end
        end
    end

end
                                           
% find all ICs with bumps btw 5 and 15 Hz                                                                 
% loop through artifacted ICs and remove any ICs that have bumps     
remove_indices=[];                                                    
for ii = 1:length(horizontal)                                         
    for jj = 1:length(cwb)%components_with_bumps)                     
        if horizontal(ii) == cwb(jj)%components_with_bumps(jj)        
            remove_indices = [remove_indices ii];                     
        end                                                           
    end                                                               
end                                                                   
horizontal(remove_indices)=[];                                        

% makes it so the func works for 64 and 128 chan nets           
if length(horizontal) > 0 %if there are artifacts                
    % eye_electrodes is entered into the func below that checks 
    % that the eye artifacts doesn't go too far into the head   
    eye_electrodes=[];                                          
    for e=1:n                                                   
        if(chanlocs(1,e).radius>0.51 && chanlocs(1,e).radius<0.60) 
            if((abs(chanlocs(1,e).theta))>27 && abs(chanlocs(1,e).theta)<58)
                eye_electrodes(e) = e;                          
            end                                                 
        end                                                     
    end                                                         
    eye_electrodes = nonzeros(eye_electrodes);                  
    sec = Spatial_Info_eyes(horizontal,eye_electrodes,topog,chanlocs,n);

% Spatial_Info_eyes(...) function is copy and pasted below for reference

function ICs_to_keep = Spatial_Info_eyes(blinks,eye_electrodes,topog,chanlocs,n)
% function discs_to_keep = Spatial_Info_discs(EEG)
%
% figures out if eye stuff rejected by adjust should be kept
%    - keeps them if there's too much spread

%define variables
ICs_to_keep = [];
arb_dist = 5.5;
arb_zval = 2;

% Get Z-Scores Matrix
zmatrix = zeros(n,n);
for i=1:n % for each topography 
    curr_vect = zeros(n);
    %deep copy ICA matrix
    for h=1:n
        zmatrix(i,h) = topog(i,h);
    end
    for h=1:n
        curr_vect(h,1) = topog(i,h);
    end
    z_vect = zscore(curr_vect);
    for h=1:n
        zmatrix(i,h) = z_vect(h,1);
    end    
end                                               

%list of channels on the outside of the head
% Find electrodes in outer ring
dimouterring=0; %number of electrodes
indexor=zeros(1,n); %indices of electrodes
for k=1:n
   if (chanlocs(1,k).radius>0.51)
        dimouterring=dimouterring+1; %count electrodes
        indexor(1,dimouterring)=k;
   end
end
outer_ring = nonzeros(indexor);
% 64-chan net: outer_ring = [1 17 18 23 24 29 32 43 47 52 55 58];
% 128-chan net: outer_ring = [48 43 49 56 63 68 73 81 88 94 99 107 113 120 119 17 128 125 127 126];
outer_ring = unique([outer_ring' eye_electrodes']);

dimhigherthresh=0;
indexht=zeros(1,n);
for k=1:n
    if (chanlocs(1,k).radius>0.35 && chanlocs(1,k).radius<0.5 && abs(chanlocs(1,k).theta)<20)
        dimhigherthresh=dimhigherthresh+1;
        indexht(1,dimhigherthresh)=k;
    end
end
higherthresh = nonzeros(indexht)';

keep_IC = []; % will be the list of ICs to keep
for ic=1:n
    if sum(ic==blinks)>0 % if this IC matches any numbers in the eye artifact list
        % find any channels/electrodes that have high activity (z>2)
        chans=find(abs(zmatrix(ic,:))>arb_zval); %grabs any chans/electrodes that are above thresh
        
        % delete the outer ring(s) of electrodes
        % We only care about the spread over more central sites for eye stuff
        chans_to_delete=[];
        for e=1:length(chans)
            if find(outer_ring == chans(e)) > 0
                % delete this chan/electrode from the list of chans (we don't care about it)
                chans_to_delete=[chans_to_delete e];
            end
            if find(higherthresh == chans(e)) > 0 
                if zmatrix(ic,chans(e))<2.5
                    % this channel isn't above the higher threshold set for this ring on the net
                    chans_to_delete=[chans_to_delete e];
                end
            end
        end
        chans(unique(chans_to_delete))=[];
        
        % calculate the distance between the points
        pts_matrix_3D=[];
        dist_matrix=[];
        for c=1:length(chans) % loop through list of chans above the threshold value and get a list of points
            pts_matrix_3D=[pts_matrix_3D; chanlocs(1,chans(c)).X chanlocs(1,chans(c)).Y chanlocs(1,chans(c)).Z];
        end
        dist_matrix = pdist(pts_matrix_3D,'euclidean'); % calculate distances btw all pts. above the threshold
        
        % figure out if component should be kept
        corresponding_pts = []; % gives us the indices of the points
        for jj=1:length(chans)-1
            corresponding_pts = [corresponding_pts; (jj*ones(1,length(chans)-jj))' ((jj+1):length(chans))'];
        end
        dists_below_thresh = find(dist_matrix<arb_dist); % tells us distances below the threshold
        pts_within_dist = corresponding_pts(dists_below_thresh,:); % tells us which point pairs are within the distnace threshold
        within_diff = [];
        if length(pts_within_dist)>0 % if any pts are within the arbitrary distance
            most_com = mode(pts_within_dist(:)); % lets us know if eye stuff spreads too far into the head
            if length(find(pts_within_dist(:) == most_com))>1
                keep_IC = [keep_IC ic];
            end
        end
        ICs_to_keep = unique(keep_IC);
    else
        % skip (we don't need to waste time looking at it if it won't be rejected)
    end
end

end % end of function

% end Spatial_Info_eyes(...) function
%%%%%%%%%%%%%%%%%%%%%%%%%%%%%%%%%%%%%%%%%%%%%%

    remove_indices=[];                                          
    for ii = 1:length(horizontal)                               
        for jj = 1:length(sec)%horiz_to_keep        )           
            if horizontal(ii) == sec(jj)                        
                remove_indices = [remove_indices ii];           
            end                                                 
        end                                                     
    end                                                         
    horizontal(remove_indices)=[];                              
end                                                             

disp('End Horizontal Eye Movement Detection');

% end of beall_horizontal(...) function
%%%%%%%%%%%%%%%%%%%%%%%%%%%%%%%%%%%%%%%%%%%%%%%%%%%%%%%%%%%%%%%%%%%%%%%%%

% intersect(intersect(find(SED>=soglia_SED),find(medie_left.*medie_right<0)),...
%     (find(nuovaV>=soglia_V)));
%horiz = intersect(find(nuovaV>=soglia_V),horiz)

hor_bool=1; %true if there are artifacted ICs

if isempty(horiz) %no IC found
    
    fprintf(file,'/ \n');
    
    hor_bool=0;
    
else
    
    fprintf(file,[num2str(horiz) '\n']);
    fprintf(file,'\n');
    
end


%% Vertical eye movements (VEM)

disp('Vertical Eye Movements...')

% ----------------------------------------------------
% |  Writes VEM header in the report file            |
% ----------------------------------------------------


fprintf(file,'>> VEM - Vertical movements\n\n');

fprintf(file,'Classification based on features:\n');

fprintf(file,'SAD - Spatial average difference (threshold=%f)\n',soglia_SAD);

fprintf(file,'MEV - Maximum epoch variance (threshold=%f)\n\n',soglia_V);

fprintf(file,'ICs with Vertical eye movements:\n');


vert=intersect(intersect(find(SAD>=soglia_SAD),find(medie_left.*medie_right>0)),...
    intersect(find(diff_var>0),find(nuovaV>=soglia_V)));
        


ver_bool=1; %true if there are artifacted ICs
        
if isempty(vert) %no artifact found
    
    fprintf(file,'/ \n');
    
    ver_bool=0;
else
    
    fprintf(file,[num2str(vert) '\n']);
    fprintf(file,'\n');    
end


%% Eye Blink (EB)

disp('Eye Blinks...')

% ----------------------------------------------------
% |  Writes EB header in the report file             |
% ----------------------------------------------------

fprintf(file,'>>> EB - Blinks\n\n');

fprintf(file,'Classification based on features:\n');

fprintf(file,'SAD (threshold=%f)\n',soglia_SAD);

fprintf(file,'TK - Temporal kurtosis (threshold=%f)\n\n',soglia_K);

fprintf(file,'ICs with Blinks:\n');

%%%%%%%%%%%%%%%%%%%%%%%%%%%%%%%%%%%%%%%%%%%%%%%%%%%%%%%%%%%%%%%%%%%%%%%%%
% adjusted_ADJUST modification                                        %%%
% blink detection code                                                %%%
blink=beall_blink_detection(cwb,topografie,EEG.chanlocs(1,pos_channels),size(EEG.icawinv,2));

% beall_blink_detection(...) function is copy and pasted below for reference

% Author: Daniel J. Beall
%
%
% beall_blink_detection() - Returns list of ICA components marked as a blink 
%
% Usage:
%   >> [blinks]=beall_blink_detection(cwb,topog,chanlocs,n);
%
% Inputs:
%   cwb        - list of components with bumps
%   topog      - topographies vector
%   chanlocs   - EEG.chanlocs struct
%   n          - number of ICs
%   nchannels  - number of channels
% Outputs:
%   blinks     - list of detected blinks
%

function [blinks] = beall_blink_detection(cwb,topog,chanlocs,n)

blinks = [];
nchannels=length(chanlocs);

%% Get Z-Scores Matrix
zmatrix = zeros(n,n);
for i=1:n % for each topography 
    curr_vect = zeros(n);
    %deep copy ICA matrix
    for h=1:n
        zmatrix(i,h) = topog(i,h);
    end
    for h=1:n
        curr_vect(h,1) = topog(i,h);
    end
    z_vect = zscore(curr_vect);
    for h=1:n
        zmatrix(i,h) = z_vect(h,1);
    end    
end
%            _                       _
% zmatrix = | z11    z12    z13   ... | row1 = IC1
%           | z21    z22    z23   ... | row2 = IC2
%           | z31    z32    z33   ... | ...
%           |_...    ...    ...   zij_|
%          col1=E1  col2=E2 ...             nxn matrix (n=#components)

%% Define scalp zones

% Find electrodes in Frontal Area (FA)
dimlefteyes=0; %number of electrodes
index1=zeros(1,nchannels); %indexes of electrodes
for k=1:nchannels
   if (chanlocs(1,k).theta > -60 && chanlocs(1,k).theta < 0) && (chanlocs(1,k).radius>0.45) && (chanlocs(1,k).radius<0.60)
        dimlefteyes=dimlefteyes+1; %count electrodes
        index1(1,dimlefteyes)=k;
    end
end

dimrighteyes=0; %number of electrodes
index2=zeros(1,nchannels); %indexes of electrodes
for k=1:nchannels
   if (chanlocs(1,k).theta < 60 && chanlocs(1,k).theta > 0) && (chanlocs(1,k).radius>0.45) && (chanlocs(1,k).radius<0.60)
        dimrighteyes=dimrighteyes+1; %count electrodes
        index2(1,dimrighteyes)=k;
    end
end

dimcentereyes=0; %number of electrodes
index3=zeros(1,nchannels); %indexes of electrodes
for k=1:nchannels
    if(abs(chanlocs(1,k).theta) < 20) && (chanlocs(1,k).radius>0.45) %electrodes are in FA
        dimcentereyes=dimcentereyes+1; %count electrodes
        index3(1,dimcentereyes)=k;
    end
end
%add in code in case E5 and E10 are deleted in a 64-chan net
if dimcentereyes==0
    for k=1:nchannels
        if(abs(chanlocs(1,k).theta) == 0) && (chanlocs(1,k).radius>0.39)
            dimcentereyes=dimcentereyes+1; %count electrodes
            index3(1,dimcentereyes)=k;
        end
    end
end

% Find electrodes in everywhere else
dimcenter=0; %number of electrodes
indexc=zeros(1,nchannels); %indexes of electrodes
for k=1:nchannels
    if( (abs(chanlocs(1,k).theta)) > 35 && abs(chanlocs(1,k).theta) < 109 && (chanlocs(1,k).radius<0.45) )
        dimcenter=dimcenter+1; %count electrodes
        indexc(1,dimcenter)=k;
    end
end

dimbackleft=0; %number of electrodes
indexbl=zeros(1,nchannels); %indexes of electrodes
for k=1:nchannels
   if (chanlocs(1,k).theta <= -109) && (chanlocs(1,k).radius<0.55) %electrodes are in FA
        dimbackleft=dimbackleft+1; %count electrodes
        indexbl(1,dimbackleft)=k;
    end
end

dimbackright=0; %number of electrodes
indexbr=zeros(1,nchannels); %indexes of electrodes
for k=1:nchannels
   if (chanlocs(1,k).theta >= 109) && (chanlocs(1,k).radius<0.55) %electrodes are in FA
        dimbackright=dimbackright+1; %count electrodes
        indexbr(1,dimbackright)=k;
    end
end

%% z-scores of the front and back

% NOTE: 
% ica struct is used for debugging purposed only
% and has no impact on the actual script run,
% just ignore it.

blinksCount = 1;
for i=1:n % for each topography
    
    % create FA electrodes vector
    zfrontlefteyes=zeros(1,dimlefteyes);
    for h=1:dimlefteyes
        zfrontlefteyes(1,h)=zmatrix(i,index1(1,h));
    end
    ica(i).lefteyeArr = zfrontlefteyes;
    
    zfrontrighteyes=zeros(1,dimrighteyes);
    for h=1:dimrighteyes
        zfrontrighteyes(1,h)=zmatrix(i,index2(1,h));
    end
    ica(i).righteyeArr = zfrontrighteyes;
    
    zcentereyes=zeros(1,dimcentereyes);
    for h=1:dimcentereyes
        zcentereyes(1,h)=zmatrix(i,index3(1,h));
    end
    ica(i).centereyeArr = zcentereyes;
    
    % create other electrodes vectors
    zcenter=zeros(1,dimcenter);
    for h=1:dimcenter
        zcenter(1,h)=zmatrix(i,indexc(1,h));
    end
    ica(i).chArr = zcenter;
    
    zbackl=zeros(1,dimbackleft);
    for h=1:dimbackleft
        zbackl(1,h)=zmatrix(i,indexbl(1,h));
    end
    ica(i).blArr = zbackl;

    zbackr=zeros(1,dimbackright);
    for h=1:dimbackright
        zbackr(1,h)=zmatrix(i,indexbr(1,h));
    end
    ica(i).brArr = zbackr;
   
    % Calculates the mean of each z-score vector
    % Sums the absolute value of each element in the vector,
    % then divides that by the number of elements.
    le = 0; 
    for j=1:dimlefteyes
        le = le + abs(zfrontlefteyes(1,j)); 
    end
    le = le/dimlefteyes;
    ica(i).lemean = le;
    
    re = 0; 
    for j=1:dimrighteyes
        re = re + abs(zfrontrighteyes(1,j)); 
    end
    re = re/dimrighteyes;
    ica(i).remean = re;
    
    ce = 0; 
    for j=1:dimcentereyes
        ce = ce + zcentereyes(1,j); %removed abs() around zcentereyes b/c they should be the same sign (1/29/19)
    end
    ce = ce/dimcentereyes;
    ica(i).cemean = ce;

    ch = 0; 
    for j=1:dimcenter
        ch = ch + abs(zcenter(1,j)); 
    end
    ch = ch/dimcenter;
    ica(i).chmean = ch;

    bl = 0; 
    for j=1:dimbackleft
        bl = bl + abs(zbackl(1,j)); 
    end
    bl = bl/dimbackleft;
    ica(i).blmean = bl;

    br = 0; 
    for j=1:dimbackright
        br = br + abs(zbackr(1,j)); 
    end
    br = br/dimbackright;
    ica(i).brmean = br;

    % if one of the means of the eye vectors pass a certain threshold and 
    % all of the back are below a certain threshold, then we reject
    if (abs(le) > 2 || abs(ce) > 2.5 || abs(re) > 2) %if high activity at front
        if (mean([abs(bl) abs(br) abs(ch)]) < 1) % if low activity in back
            if ( (var([zbackl zbackr]') < 0.15 || var([abs(zbackl) abs(zbackr)]')<0.075) && var([zfrontlefteyes zfrontrighteyes]')>var([zbackl zbackr]') ) %if low variance in back and greater variance in front
                blinks(blinksCount) = i;
                blinksCount = blinksCount + 1;
            end
        end
    end

end
                                        
%find all ICs with bumps btw 5 and 15 Hz                                                                   
%loop through artifacted ICs and remove any ICs that have bumps       
remove_indices=[];                                                    
for ii = 1:length(blinks)                                             
    for jj = 1:length(cwb)%components_with_bumps)                     
        if blinks(ii) == cwb(jj)%components_with_bumps(jj)            
            remove_indices = [remove_indices ii];                    
        end                                                           
    end                                                              
end                                                                   
blinks(remove_indices)=[];                                           

                           
if length(blinks)>0                                            
    % eye_electrodes is entered into the func below to check   
    % that the eye artifacts doesn't go too far into the head  
    eye_electrodes=[];                                         
    for e=1:n                                                  
        if(chanlocs(1,e).radius>0.45 && chanlocs(1,e).radius<0.54) 
            if(abs(chanlocs(1,e).theta)<17)                   
                eye_electrodes(e) = e;                         
            end                                                
        end                                                    
    end                                                        
    eye_electrodes = nonzeros(eye_electrodes);                 
    sec = Spatial_Info_eyes(blinks,eye_electrodes,topog,chanlocs,n); 

% Spatial_Info_eyes(...) function is copy and pasted below for reference

function ICs_to_keep = Spatial_Info_eyes(blinks,eye_electrodes,topog,chanlocs,n)
% function discs_to_keep = Spatial_Info_discs(EEG)
%
% figures out if eye stuff rejected by adjust should be kept
%    - keeps them if there's too much spread

%define variables
ICs_to_keep = [];
arb_dist = 5.5;
arb_zval = 2;

% Get Z-Scores Matrix
zmatrix = zeros(n,n);
for i=1:n % for each topography 
    curr_vect = zeros(n);
    %deep copy ICA matrix
    for h=1:n
        zmatrix(i,h) = topog(i,h);
    end
    for h=1:n
        curr_vect(h,1) = topog(i,h);
    end
    z_vect = zscore(curr_vect);
    for h=1:n
        zmatrix(i,h) = z_vect(h,1);
    end    
end                                               

%list of channels on the outside of the head
% Find electrodes in outer ring
dimouterring=0; %number of electrodes
indexor=zeros(1,n); %indices of electrodes
for k=1:n
   if (chanlocs(1,k).radius>0.51)
        dimouterring=dimouterring+1; %count electrodes
        indexor(1,dimouterring)=k;
   end
end
outer_ring = nonzeros(indexor);
% 64-chan net: outer_ring = [1 17 18 23 24 29 32 43 47 52 55 58];
% 128-chan net: outer_ring = [48 43 49 56 63 68 73 81 88 94 99 107 113 120 119 17 128 125 127 126];
outer_ring = unique([outer_ring' eye_electrodes']);

dimhigherthresh=0;
indexht=zeros(1,n);
for k=1:n
    if (chanlocs(1,k).radius>0.35 && chanlocs(1,k).radius<0.5 && abs(chanlocs(1,k).theta)<20)
        dimhigherthresh=dimhigherthresh+1;
        indexht(1,dimhigherthresh)=k;
    end
end
higherthresh = nonzeros(indexht)';

keep_IC = []; % will be the list of ICs to keep
for ic=1:n
    if sum(ic==blinks)>0 % if this IC matches any numbers in the eye artifact list
        % find any channels/electrodes that have high activity (z>2)
        chans=find(abs(zmatrix(ic,:))>arb_zval); %grabs any chans/electrodes that are above thresh
        
        % delete the outer ring(s) of electrodes
        % We only care about the spread over more central sites for eye stuff
        chans_to_delete=[];
        for e=1:length(chans)
            if find(outer_ring == chans(e)) > 0
                % delete this chan/electrode from the list of chans (we don't care about it)
                chans_to_delete=[chans_to_delete e];
            end
            if find(higherthresh == chans(e)) > 0 
                if zmatrix(ic,chans(e))<2.5
                    % this channel isn't above the higher threshold set for this ring on the net
                    chans_to_delete=[chans_to_delete e];
                end
            end
        end
        chans(unique(chans_to_delete))=[];
        
        % calculate the distance between the points
        pts_matrix_3D=[];
        dist_matrix=[];
        for c=1:length(chans) % loop through list of chans above the threshold value and get a list of points
            pts_matrix_3D=[pts_matrix_3D; chanlocs(1,chans(c)).X chanlocs(1,chans(c)).Y chanlocs(1,chans(c)).Z];
        end
        dist_matrix = pdist(pts_matrix_3D,'euclidean'); % calculate distances btw all pts. above the threshold
        
        % figure out if component should be kept
        corresponding_pts = []; % gives us the indices of the points
        for jj=1:length(chans)-1
            corresponding_pts = [corresponding_pts; (jj*ones(1,length(chans)-jj))' ((jj+1):length(chans))'];
        end
        dists_below_thresh = find(dist_matrix<arb_dist); % tells us distances below the threshold
        pts_within_dist = corresponding_pts(dists_below_thresh,:); % tells us which point pairs are within the distnace threshold
        within_diff = [];
        if length(pts_within_dist)>0 % if any pts are within the arbitrary distance
            most_com = mode(pts_within_dist(:)); % lets us know if eye stuff spreads too far into the head
            if length(find(pts_within_dist(:) == most_com))>1
                keep_IC = [keep_IC ic];
            end
        end
        ICs_to_keep = unique(keep_IC);
    else
        % skip (we don't need to waste time looking at it if it won't be rejected)
    end
end

end % end of function

% end Spatial_Info_eyes(...) function
%%%%%%%%%%%%%%%%%%%%%%%%%%%%%%%%%%%%%%%%%%%%%%

    remove_indices=[];                                         
    for ii = 1:length(blinks)                                  
        for jj = 1:length(sec)%blinks_to_keep                  
            if blinks(ii) == sec(jj)                           
                remove_indices = [remove_indices ii];          
            end                                                
        end                                                    
    end                                                       
    blinks(remove_indices)=[];                                 
end                                                            

disp('End Blink Detection');

% end beall_blink_detection(...) function
%%%%%%%%%%%%%%%%%%%%%%%%%%%%%%%%%%%%%%%%%%%%%%%%%%%%%%%%%%%%%%%%%%%%%%%%%%

% intersect ( intersect( find(SAD>=soglia_SAD),find(medie_left.*medie_right>0) ) ,...
%     intersect ( find(meanK>=soglia_K),find(diff_var>0) ));
% blink = intersect( find(meanK>=soglia_K), blink )

bl_bool=1; %true if there are artifacted ICs
            
if isempty(blink) %no blink component
    
    fprintf(file,'/ \n');
    
    bl_bool=0;
else
    
    fprintf(file,[num2str(blink) '\n']);
    fprintf(file,'\n');    
end


%% Generic Discontinuities (GD)

disp('Generic Discontinuities...')

% ----------------------------------------------------
% |  Writes GD header in the report file             |
% ----------------------------------------------------

fprintf(file,'>>>> GD - Discontinuities\n');

fprintf(file,'Classification based on features:\n');

fprintf(file,'GDSF - Generic Discontinuities Spatial Feature (threshold=%f)\n',soglia_GDSF);

fprintf(file,'MEV - Maximum epoch variance (threshold=%f)\n\n',soglia_V);

fprintf(file,'ICs with Generic Discontinuities:\n');


disc=intersect(find(GDSF>=soglia_GDSF),find(nuovaV>=soglia_V));


dsc_bool=1; %true if there are discontinuities
               
if isempty(disc) %no discontinuities
    
    fprintf(file,'/ \n');
    
    dsc_bool=0;
else
    
    fprintf(file,[num2str(disc) '\n']);
    fprintf(file,'\n');    
end

%%

aic=unique([blink disc horiz vert]);
%%%%%%%%%%%%%%%%%%%%%%%%%%%%%%%%%%%%%%%%%%%%%%%%%%%%%%%%%%%%%%%%%%%%%%
% adjusted_ADJUST modification                                        %
%loop through artifacted ICs and remove any ICs that have bumps       %
remove_indices=[];                                                    %
for ii = 1:length(aic)                                                %
    for jj = 1:length(cwb) %components_with_bumps                     %
        if aic(ii) == cwb(jj)%components_with_bumps(jj)               %
            remove_indices = [remove_indices ii];                     %
        end                                                           %
    end                                                               %
end                                                                   %
aic(remove_indices)=[];                                               %
%%%%%%%%%%%%%%%%%%%%%%%%%%%%%%%%%%%%%%%%%%%%%%%%%%%%%%%%%%%%%%%%%%%%%%

fprintf(file,'Artifacted ICs (total):\n');
    fprintf(file,[num2str(aic) '\n']);
    fprintf(file,'\n');    


%% Displaying results

% ----------------------------------------------------
% |  Write message to user: report file name         |
% ----------------------------------------------------

disp(' ')
disp(['Results in <' out '>.'])


fclose(file);

%compute output variable
art = nonzeros( union (union(blink,horiz) , union(vert,disc)) )'; %artifact ICs
%%%%%%%%%%%%%%%%%%%%%%%%%%%%%%%%%%%%%%%%%%%%%%%%%%%%%%%%%%%%%%%%%%%%%%
% adjusted_ADJUST modification                                        %
remove_indices=[];                                                    %
for ii = 1:length(art)                                                %
    for jj = 1:length(cwb)%components_with_bumps                      %
        if art(ii) == cwb(jj)%components_with_bumps(jj)               %
            remove_indices = [remove_indices ii];                     %
        end                                                           %
    end                                                               %
end                                                                   %
art(remove_indices)=[];                                               %
%%%%%%%%%%%%%%%%%%%%%%%%%%%%%%%%%%%%%%%%%%%%%%%%%%%%%%%%%%%%%%%%%%%%%%

% these three are old outputs which are no more necessary in latest ADJUST version.
soglia_D=0;
soglia_DV=0;
maxdin=zeros(1,size(EEG.icawinv,2));

return

%% The following sections have been moved to interface_ADJ in order to manage
%% continuous data

% 
% %% Saving artifacted ICs for further analysis
% 
% nome=['List_' EEG.setname '.mat'];
% 
% save (nome, 'blink', 'horiz', 'vert', 'disc');
% 
% disp(' ')
% disp(['Artifact ICs list saved in ' nome]);
% 
% 
% %% IC show & remove
% % show all ICs; detected ICs are highlighted in red color. Based on
% % pop_selectcomps.
% 
% art = nonzeros( union (union(blink,horiz) , union(vert,disc)) )'; %artifact ICs
% 
% %     [EEG] = pop_selectcomps_ADJ( EEG, 1:size(EEG.icawinv,1), art, horiz, vert, blink, disc,...
% %         soglia_DV, diff_var, soglia_K, meanK, soglia_SED, SED, soglia_SAD, SAD, ...
% %         soglia_TDR, topog_DR, soglia_V, maxvar, soglia_D, maxdin );
%     [EEG] = pop_selectcomps_ADJ( EEG, 1:size(EEG.icawinv,1), art, horiz, vert, blink, disc,...
%         soglia_DV, diff_var, soglia_K, med2_K, meanK, soglia_SED, med2_SED, SED, soglia_SAD, med2_SAD, SAD, ...
%         soglia_GDSF, med2_GDSF, topog_DR, soglia_V, med2_V, maxvar, soglia_D, maxdin );
